# Competition for electrons favors N_2_O reduction in denitrifying *Bradyrhizobium isolates*

**DOI:** 10.1101/2020.07.20.212696

**Authors:** Y Gao, D Mania, SA Mousavi, P Lycus, M Arntzen, K Woliy, K Lindström, JP Shapleigh, LR Bakken, Å Frostegård

**Affiliations:** Faculty of Chemistry, Biotechnology and Food Sciences, Norwegian University of Life Sciences, Aas, Norway; Ecosystems and Environment Research programme, Faculty of Biological and Environmental Sciences, and Helsinki Institute of Sustainability Science (HELSUS), University of Helsinki, Finland; Department of Microbiology, Cornell University, Ithaca, New York, USA

## Abstract

Bradyrhizobia are common members of soil microbiomes and known as N_2_-fixing symbionts of economically important legumes. Many are also denitrifiers, which can act as sinks or sources for N_2_O. Inoculation with compatible rhizobia is often needed for optimal N_2_-fixation, but the choice of inoculant may also have consequences for N_2_O emission. Here, we analyzed the phylogeny and denitrification capacity of *Bradyrhizobium* strains, most of them isolated from peanut-nodules. All were dinitrifiers, but only ^~^1/3 could reduce N_2_O while most others were net N_2_O producers. The N_2_O-reducing isolates showed strong preference for N_2_O- over NO_3_^−^-reduction. Such preference was also observed in a study of other bradyrhizobia and tentatively ascribed to competition between the electron pathways to Nap (periplasmic NO_3_^−^ reductase) and Nos (N_2_O reductase). Another possible explanation is lower abundance of Nap than Nos. Here, proteomics revealed that Nap was instead more abundant than Nos, supporting the hypothesis that the electron pathway to Nos outcompetes that to Nap. In contrast, *Paracoccus denitrificans, which has membrane-bond* NO_3_^−^ reductase (Nar), reduced N_2_O and NO_3_^−^ simultaneously. We propose that the control at the metabolic level, favoring N_2_O reduction over NO_3_^−^ reduction, applies also to other denitrifiers carrying Nos and Nap but lacking Nar.

**Originality-Significance Statement:** This study extends the current knowledge on denitrification in bradyrhizobia, which mostly originates from studies of one model strain, by investigating the denitrification phenotypes of a diverse collection of Bradyrhizobium isolates. Only 1/3 of them could reduce N_2_O while the others were net sources for this potent greenhouse gas. All N_2_O-reducers showed strong preference for N_2_O over NO_3_^−^. We revealed by proteomics that this was not explained by differences in the abundances of Nap (periplasmic nitrate reductase) and Nos (N_2_O reductase), which strengthens our hypothesis (Mania *et al*., 2020) of a metabolic control mechanism by which Nos competes efficiently with Nap for electrons, making these organisms strong sinks for N_2_O. The findings highlight the potential importance of these organisms as N_2_O sinks in natural and agricultural ecosystems and pinpoint the need to take N_2_O reduction into account, along with N_2_-fixation effectiveness, when searching for strains suitable for production of inoculants.

## Introduction

Emissions of nitrous oxide (N_2_O), the third most important anthropogenic greenhouse gas, have been rising steadily during the past 150 years. Predictions for the rest of the century show either continued N_2_O emission increases or rates of decrease slower than the decreases occurring in CO_2_ emissions (Rogelj *et al*., 2018; Thompson *et al.*, 2019). Agriculture is the main source of anthropogenic N_2_O emissions, with denitrification playing the major role in most soils (Thomson *et al.*, 2012). The primary cause of the emissions is excessive use of synthetic fertilizers (Reay *et al.*, 2012; Thompson *et al.*, 2019; Tian *et al.*, 2020), but cultivation of legumes also contributes substantially with a global average of 1.29 (range 0.03-7.09) kg N_2_O-N ha^−1^ year^−1^ while the corresponding estimate for fertilized agricultural fields was 3.22 (range 0.09-12.67) (Jensen *et al.*, 2012). In both cases, the ultimate driver of the emission is input of reactive nitrogen which undergoes a series of redox reactions and eventually returns to the atmosphere as N_2_O or N_2_. Our aim should be to minimize the N_2_O/N_2_ emission ratio of the system (Schlesinger, 2009). To achieve this, novel mitigation options beyond “good agronomic practice” will be needed (Davidson and Kanter, 2014). Several technical abatement options have been suggested, including precision fertilization and use of nitrification inhibitors (Winiwarter *et al.*, 2018).

An alternative strategy is to enhance the populations of N_2_O reducing microorganisms in the soil. Selective *in situ* stimulation of N_2_O reducing, indigenous soil organisms appears unrealistic, while some options for the addition of N_2_O reducing microbes may be feasible. One is the suggestion to inoculate soybeans with rhizobia that are not only efficient N_2_-fixers but also have the capacity to reduce N_2_O (Hénault and Revellin, 2011; Itakura *et al.*, 2013; Akiyama *et al.*, 2016). At present, N_2_O reduction is not taken into account when screening strains for their suitability as inoculants and, in fact, several of the commercial inoculants currently used are only partial denitrifiers (Zilli *et al.*, 2019). For example, only one of the five *Bradyrhizobium* strains most commonly used as soybean inoculants in South America could reduce N_2_O (Obando *et al.*, 2019). The growing global need for plant-based proteins is expected to require new areas to be used for legume cropping and will likely increase the demand for new legume varieties. Not all combinations of legume cultivar and rhizobial strain result in an efficient symbiosis (Thrall *et al.*, 2011; Woliy *et al*, 2019). Importantly, indigenous soil populations of rhizobia may not be compatible with a certain legume crop, especially if it has not been grown in that area before. Therefore, the expansion of legume cropping is creating an increasing demand for a wider variety of inoculants to ensure nodulation and optimal N_2_ fixation efficacy (Santos *et al.*, 2019). This offers a golden opportunity to mitigate N_2_O emissions, providing that the inoculants are selected among rhizobial strains that can reduce N_2_O.

The purpose of the present study was to determine how denitrification phenotypes vary across taxonomically diverse groups of bradyrhizobia, with particular focus on their capacity for N_2_O reduction, and to understand cellular mechanisms that make these organisms potential sinks for N_2_O. We determined the phylogenetic position of a set of *Bradyrhizobium* strains, obtained from different culture collections, and analyzed their capacity for denitrification. Most of the strains were isolated from nodules of peanut plants (*Arachis hypogaea*) growing in different regions of China. Peanut is the fourth most important legume crop globally, with China being the main producer (Arya *et al.*, 2016). We also included some bradyrhizobia from other parts of the world, isolated from peanut and other legumes.

Several taxonomic groups within the genus *Bradyrhizobium* have been reported to denitrify with many strains having truncated denitrification pathways lacking one or more of the steps (Sameshima-Saito *et al.,* 2004; Zilli *et al*., 2019; Mania *et al*., 2020). A complete denitrification pathway in bradyrhizobia comprises the periplasmic reductase Nap for dissimilatory NO_3_^−^ reduction, encoded as a part of the *napEDABC* operon; the Cu-containing nitrite reductase NirK encoded by *nirK* only or as a part of the *nirKV* operon, although *cd*_*1*_-type NirS nitrite reductase has also been reported (Sánchez and Minamisawa, 2018); the cytochrome *c* dependent nitric oxide reductase cNor *(norCBQD* operon); and a clade I N_2_O reductase Nos (*nosRZDYFLX* operon) (Velasco *et al.*, 2001;Mesa *et al.*, 2002; Delgado *et al.*, 2003; Velasco *et al.*, 2004; Bedmar *et al.*, 2005). While the bradyrhizobia, along with several other denitrifiers, only have the periplasmic Nap for dissimilatory NO_3_^−^ reduction, other denitrifiers have the membrane-bound Nar, or both. Nap is a periplasmic heterodimer (NapAB) consisting of the small electron transfer subunit NapB and the catalytic NapA, which draws electrons from quinol via the membrane-bound *c*-type tetrahaeme cytochrome NapC. Nar is a three-subunit enzyme (NarGHI) anchored in the membrane by NarI, which is a transmembrane protein that draws electrons from quinol to the catalytic NarG via NarH, both facing the cytoplasm (Richardson, 2000; Richardson *et al.*, 2001). Nir, Nor and Nos receive electrons from the *bc1* complex via periplasmic small c-type cytochromes (Berks *et al.*, 1995; Zumft, 1997). Clade I type Nos reductase may in addition draw electrons from quinol via the membrane bound NosR (Zhang *et al.*, 2019).

In a recent study, Mania *et al.* (2020) screened strains within the genus *Bradyrhizobium* for their denitrification phenotype. Most of them could perform two or more of the denitrification steps, but only half of them could reduce N_2_O. All the N_2_O reducing strains displayed the same strong preference for N_2_O over NO_3_^−^ reduction when incubated under anoxic conditions, which was suggested to be due to competition between the electron pathways to Nap and Nos. The quantification of denitrification gene transcripts showed 5-8 times lower transcript numbers of *nap* compared to *nos,* however. Therefore, it could not be excluded that the phenomenon is due to low abundance of Nap compared to Nos. Here we have investigated the denitrification geno- and phenotypes for another collection of *Bradyrhizobium* isolates and deepened our understanding of the competition for electrons between Nap and Nos by refined analyses of the electron flow kinetics, in combination with the determination of the relative amounts of Nap and Nos by proteomics. In addition, we compared the denitrification kinetics of the bradyrhizobia with the denitrification model bacterium *Paracoccus denitrificans*, which holds a complete denitrification pathway including both Nap and Nar (Zumft, 1997; Richardson, 2000). The results showed that in all bradyrhizobia with complete denitrification, NO_3_^−^ reduction was almost completely hampered in the presence of N_2_O. The proteomics analysis provided evidence that this was not due to a low abundance of Nap, also showing that transcript abundance cannot be directly translated into corresponding protein abundance. As expected, the NarG carrying *P. denitrificans* showed a different denitrification phenotype with simultaneous reduction of NO_3_^−^ and N_2_O.

## Results

### Phylogenetic distribution of denitrifying bradyrhizobia

The phylogenetic position of the 23 test strains was determined by MLSA analysis of four protein-coding housekeeping genes (*atpD-glnII-recA-rpoB*) (Fig. 1). The analysis comprised 78 bradyrhizobial strains and included all hitherto described *Bradyrhizobium* reference species. All but one of the 23 test strains listed in Table 1 were assigned to the *Bradyrhizobium japonicum* supergroup (Fig. 1). The exception was *Bradyrhizobium lablabi* HAMBI 3052^T^, which is a member of the *Bradyrhizobium jicamae* supergroup according to the new genome-based taxonomy of bradyrhizobia proposed by Avontuur *et al.* (2019). The test strains were placed in 11 lineages (clades A-K) with high posterior probability (PP=1.00). Most test strains were clustered within the clades of seven diverse bradyrhizobial species. Strains *Bradyrhizobium* sp. HAMBI 2134 and 2142 formed a monophyletic cluster (clade G) with high posterior probability (1.00). *Bradyrhizobium* sp. strain HAMBI 2116 was accommodated in a lineage (clade A) distant from other bradyrhizobia. The phylogenetic relationships revealed by the four-gene MLSA tree (Fig. 1) showed good correspondence with the results from an analysis of all the HAMBI strains using the GTDB Tool kit (Fig. S1). In both trees strains HAMBI 2299, 2153 and 2116 were most closely related to the reference species *B. yuanmingense*; strains HAMBI 2125 (and 2126) and 2127 were closest to *B. ottawaense*; strain HAMBI 2130 was closest to *B. shewense*; strains HAMBI 2115 was closest to *B. japonicum* USDA6; strains HAMBI 2128 (as well as HAMBI 2129 and 2150, Table S1) were closest to *B. arachidis*; strain HAMBI 3052 was closest to *B. lablabi.* Strains HAMBI 2149, 2133, 2136, 2137, 2151 and 2152 formed one cluster in the GTDB tree together with *B. japonicum* USDA 135. This cluster was close to *B. liaoningense*, as also found in the MLSA analysis (Clade F, Fig. 1), which did not include *B. japonicum* USDA 135. Strains HAMBI 2134 (and 2135, Table S1) and 2142 formed a separate cluster in both trees.

**Table 1.**
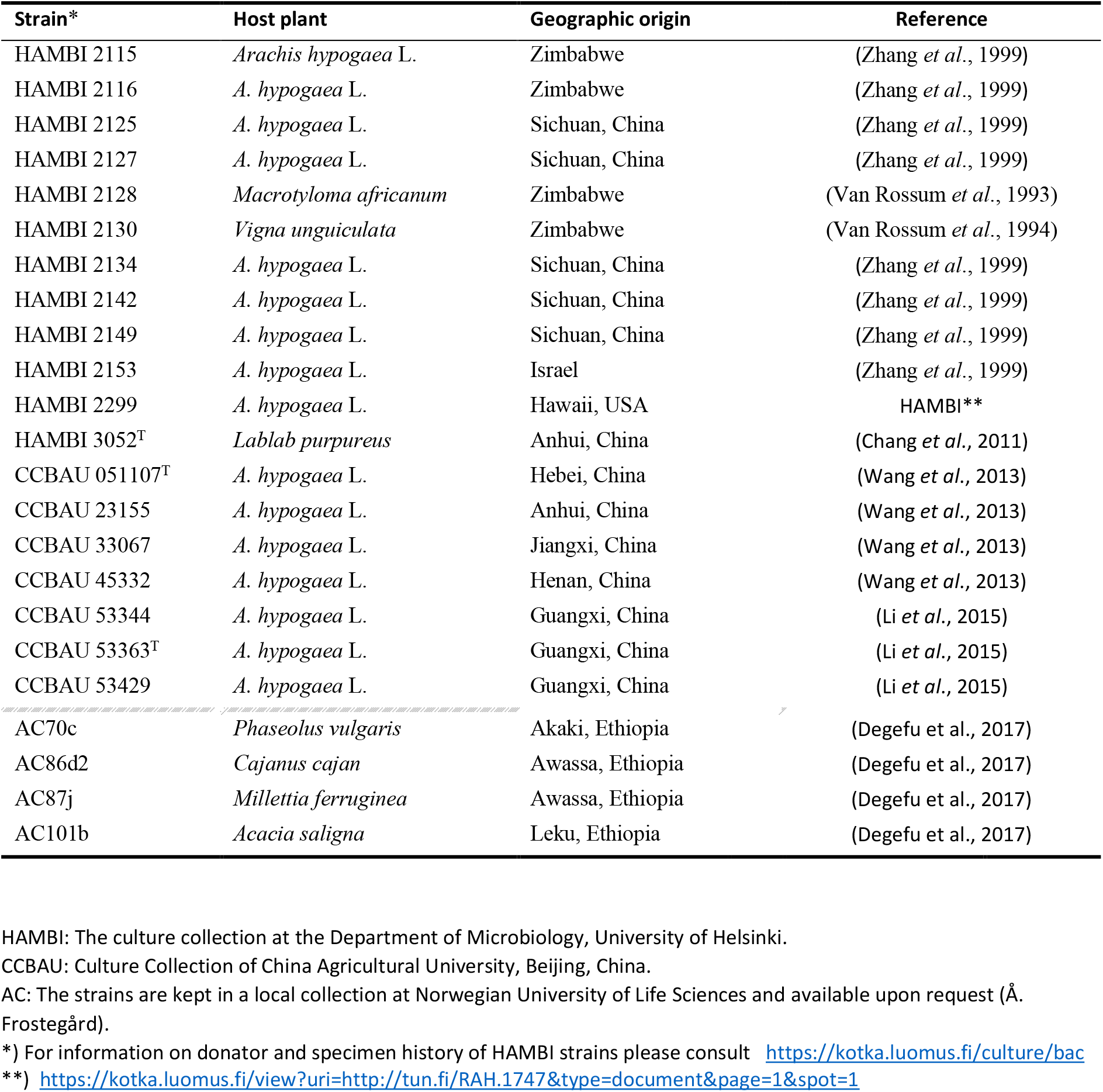
Strains used in the current study. All listed strains were included in the MLSA-based phylogenetic analysis (Fig. 1). The strains above the dashed line were also examined in the present study with regard to their denitrification phenotypes. Strains AC 70c, AC 86d2, AC 86j1 and AC 101b were not examined for denitrification since this was reported elsewhere (Mania *et al*., 2020).

| Strain* | Host plant | Geographic origin | Reference |
| --- | --- | --- | --- |
| HAMBI 2115 | *Arachis hypogaea* L. | Zimbabwe | (Zhang *et al.*, 1999) |
| HAMBI 2116 | *A. hypogaea* L. | Zimbabwe | (Zhang *et al.*, 1999) |
| HAMBI 2125 | *A. hypogaea* L. | Sichuan, China | (Zhang *et al.*, 1999) |
| HAMBI 2127 | *A. hypogaea* L. | Sichuan, China | (Zhang *et al.*, 1999) |
| HAMBI 2128 | *Macrotyloma africanum* | Zimbabwe | (Van Rossum *et al.*, 1993) |
| HAMBI 2130 | *Vigna unguiculata* | Zimbabwe | (Van Rossum *et al.*, 1994) |
| HAMBI 2134 | *A. hypogaea* L. | Sichuan, China | (Zhang *et al.*, 1999) |
| HAMBI 2142 | *A. hypogaea* L. | Sichuan, China | (Zhang *et al.*, 1999) |
| HAMBI 2149 | *A. hypogaea* L. | Sichuan, China | (Zhang *et al.*, 1999) |
| HAMBI 2153 | *A. hypogaea* L. | Israel | (Zhang *et al.*, 1999) |
| HAMBI 2299 | *A. hypogaea* L. | Hawaii, USA | HAMBI** |
| HAMBI 3052<sup>T</sup> | *Lablab purpureus* | Anhui, China | (Chang *et al.*, 2011) |
| CCBAU 051107<sup>T</sup> | *A. hypogaea* L. | Hebei, China | (Wang *et al.*, 2013) |
| CCBAU 23155 | *A. hypogaea* L. | Anhui, China | (Wang *et al.*, 2013) |
| CCBAU 33067 | *A. hypogaea* L. | Jiangxi, China | (Wang *et al.*, 2013) |
| CCBAU 45332 | *A. hypogaea* L. | Henan, China | (Wang *et al.*, 2013) |
| CCBAU 53344 | *A. hypogaea* L. | Guangxi, China | (Li *et al.*, 2015) |
| CCBAU 53363<sup>T</sup> | *A. hypogaea* L. | Guangxi, China | (Li *et al.*, 2015) |
| CCBAU 53429 | *A. hypogaea* L. | Guangxi, China | (Li *et al.*, 2015) |
| AC70c | *Phaseolus vulgaris* | Akaki, Ethiopia | (Degefu *et al.*, 2017) |
| AC86d2 | *Cajanus cajan* | Awassa, Ethiopia | (Degefu *et al.*, 2017) |
| AC87j | *Millettia ferruginea* | Awassa, Ethiopia | (Degefu *et al.*, 2017) |
| AC101b | *Acacia saligna* | Leku, Ethiopia | (Degefu *et al.*, 2017) |

HAMBI: The culture collection at the Department of Microbiology, University of Helsinki.
CCBAU: Culture Collection of China Agricultural University, Beijing, China.
AC: The strains are kept in a local collection at Norwegian University of Life Sciences and available upon request (Å. Frostegård).
\*) For information on donator and specimen history of HAMBI strains please consult https://kotka.luomus.fi/culture/bac
\*\*) https://kotka.luomus.fi/view?uri=http://tun.fi/RAH.1747&type=document&page=1&spot=1

**Fig. 1.**
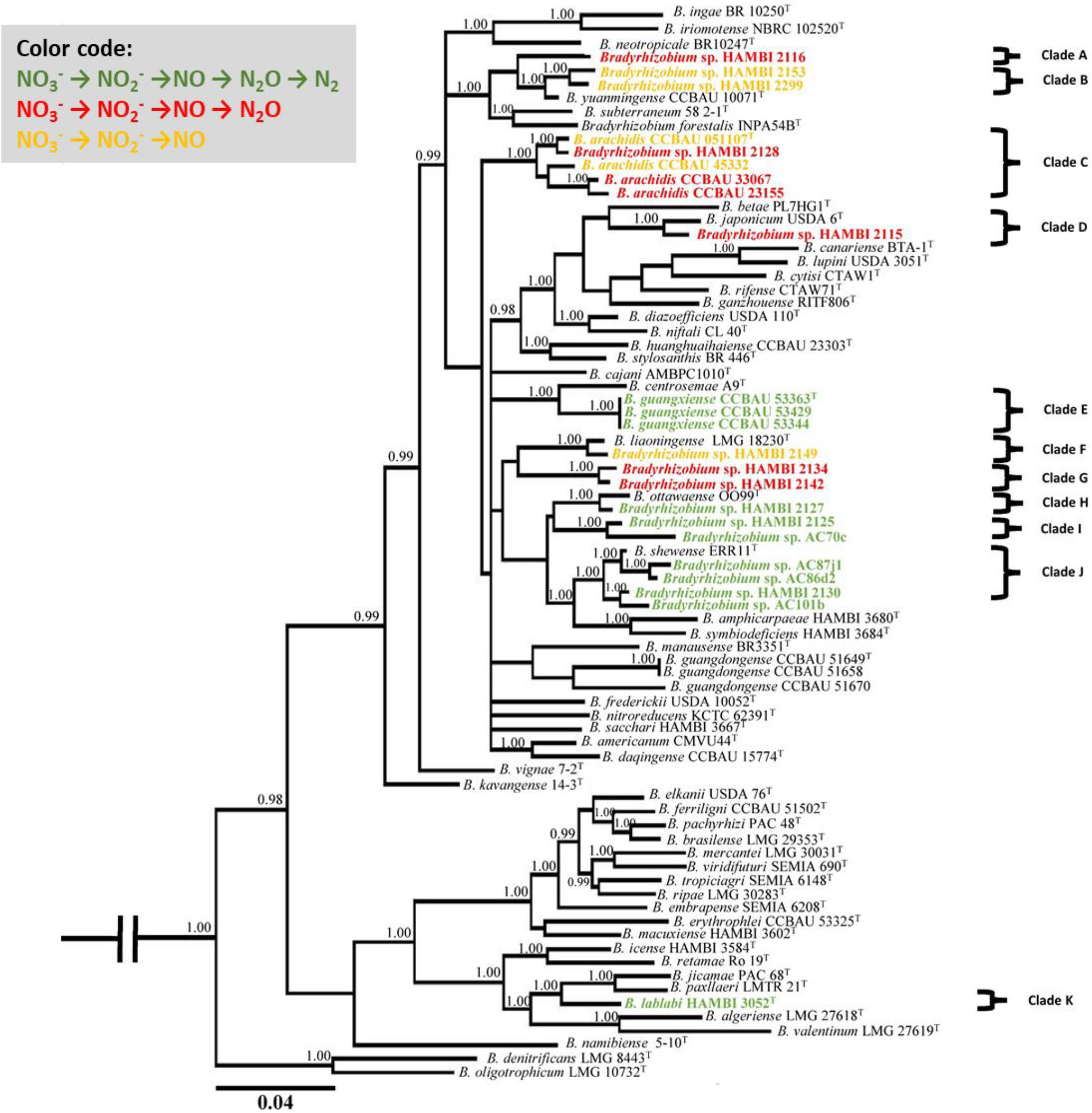
Phylogenetic tree of four concatenated housekeeping genes (*atpD-glnII-recA-rpoB*) from 78 *Bradyrhizobium* strains constructed based on a Bayesian inference analysis. Only the posterior probabilities ≥0.95 are shown in the tree. The genus name *Bradyrhizobium* is abbreviated as *B*., and a “T” at the end of each strain code shows the type strains. In cases where two or more isolates showed high whole genome sequence similarity (>99 %) based on pairwise comparison of Average Nucleotide Identity (ANI) (see Table S1), only one representative is shown in this tree. Colors indicate denitrification end-products. A sequence analysis of the genes in the respective denitrification gene operons showed good correspondence between genetic set-up and denitrification end products (Table S2). The only exception was HAMBI 2299, which had a complete *nor* operon but did not reduce NO. Another non-NO reducing strain, HAMBI 2153, lacked two genes in the *nor* operon.

We originally included 21 isolates from the HAMBI culture collection in our study, all deposited with different strain names. Pairwise whole genome comparisons revealed that some strains were very similar with ANI values >99.2% and in most cases >99.9 % (Table S1), which was also supported by the clustering of similar strains in the GTDB tree (Fig. S1). Since the strains within such clusters with ANI values >99% also showed practically identical denitrification phenotypes, we present the result for only one of the strains in each such cluster in the main paper, and the results for the others as supplementary information.

The denitrification end-product analysis showed that all test strains were denitrifiers, able to reduce NO_3_^−^ and/or NO_3_^−^ to NO, N_2_O or N_2_. Seven of the 19 strains tested for denitrification in the present study were complete denitrifers that reduced NO_3_^−^ to N_2_. The four Ethiopian strains AC87j1, AC86d2, AC101b and AC70c were determined earlier to be complete denitrifiers (Mania *et al*. 2020). All complete denitrifiers except *B. lablabi* (clade K) belonged to the *B. japonicum* supergroup and were positioned in clades E, H, I and J. The clades E, H and J represented the species *B. guangxiense*, *B. ottawaense* and *B. shewense*, respectively. Clade I, accommodating strains *Bradyrhizobium* sp. HAMBI 2125 and AC70c, formed a separate, monophyletic group. The other test strains had truncated denitrification pathways and were able to reduce NO_3_^−^ and NO_2_^−^ to NO (5 strains) or N_2_O (7 strains). The seven strains having N_2_O as end-product were distributed in clades A, C, D and G. Three strains grouped with *B. arachidis* in clade C*. Bradyrhizobium* sp. strain HAMBI 2115 was placed in clade D along with *B. japonicum* USDA6^T^. Three strains were accommodated in clades A and G, neither of which included any of the hitherto described *Bradyrhizobium* species. The strains with NO as denitrification end-product were found in clades B, C and F, which also harbored the species *B. yuanmingense*, *B. arachidis* and *B. liaoningense*, respectively. Strains HAMBI 2153 and 2149, as well as the strains highly similar to HAMBI 2149 (Table S1), were also examined using detailed gas kinetics to further examine whether they could reduce NO (Fig. S2). Like in the end-product investigation they produced large amounts of NO (14-15 μmol flask^−1^) while no reduction of NO or N_2_O was detected (Fig. S2).

### Consistency between denitrification gene operons and end products

The genetic potential for denitrification, based on whole genome sequencing, and the corresponding phenotype, determined by end-point analysis and gas kinetics, were compared for 13 strains (Table S2), as well as for isolates with genomes with ANI values >99.2% to these strains (Table S1). The five strains able to reduce NO_3_^−^ to N_2_ had complete operons for all four reduction steps. For most of the other strains, with NO or N_2_O as denitrification end -products, the lack of a function was explained by complete lack of the corresponding operon. However, two strains (HAMBI 2153 and HAMBI 2299) showed a discrepancy between denitrification genotype and phenotype. They were apparently unable to reduce NO and accumulated μM concentrations of NO, despite having the *nor* operon. Comparison of these strains with strains having functional NO reduction revealed that strain HAMBI 2153 had a frame shift in *norB*, while *norQ* was not in the correct reading frame. Strain HAMBI 2299 carried *nap, nir* and *nor* operons with all expected genes present. The end-point analysis of this strain showed that after 10 days of incubation with 1 mM NO_3_^−^, 0.5 mM NO_2_^−^ and 1 ml N_2_O in headspace, it had produced toxic levels (>10 μM) of NO. Inspection of the amino acid sequences did not reveal any obvious explanations for lack of Nor activity.

### All N_2_O-reducing Bradyrhizobium strains preferred N_2_O over NO_3_^−^ as electron acceptor

To determine if the complete denitrifiers showed a similar, strong preference for N_2_O over NO_3_^−^ as reported by Mania *et al.* (2020) for other *Bradyrhizobium* strains, detailed gas kinetics was analyzed. The results showed remarkably similar kinetics for all strains during and after the transition from aerobic to anaerobic respiration in flasks to which N_2_O had been added to the headspace (Figs 2A, S3, S4, S5). Strain HAMBI 2125 was chosen as a representative for the complete denitrifiers. The NO_2_^−^ concentration in cultures of this strain was measured along with the gas kinetics (Figs 2A, S3), allowing the estimation of NO_3_^−^ concentrations by N-mass balance.

**Fig. 2.**
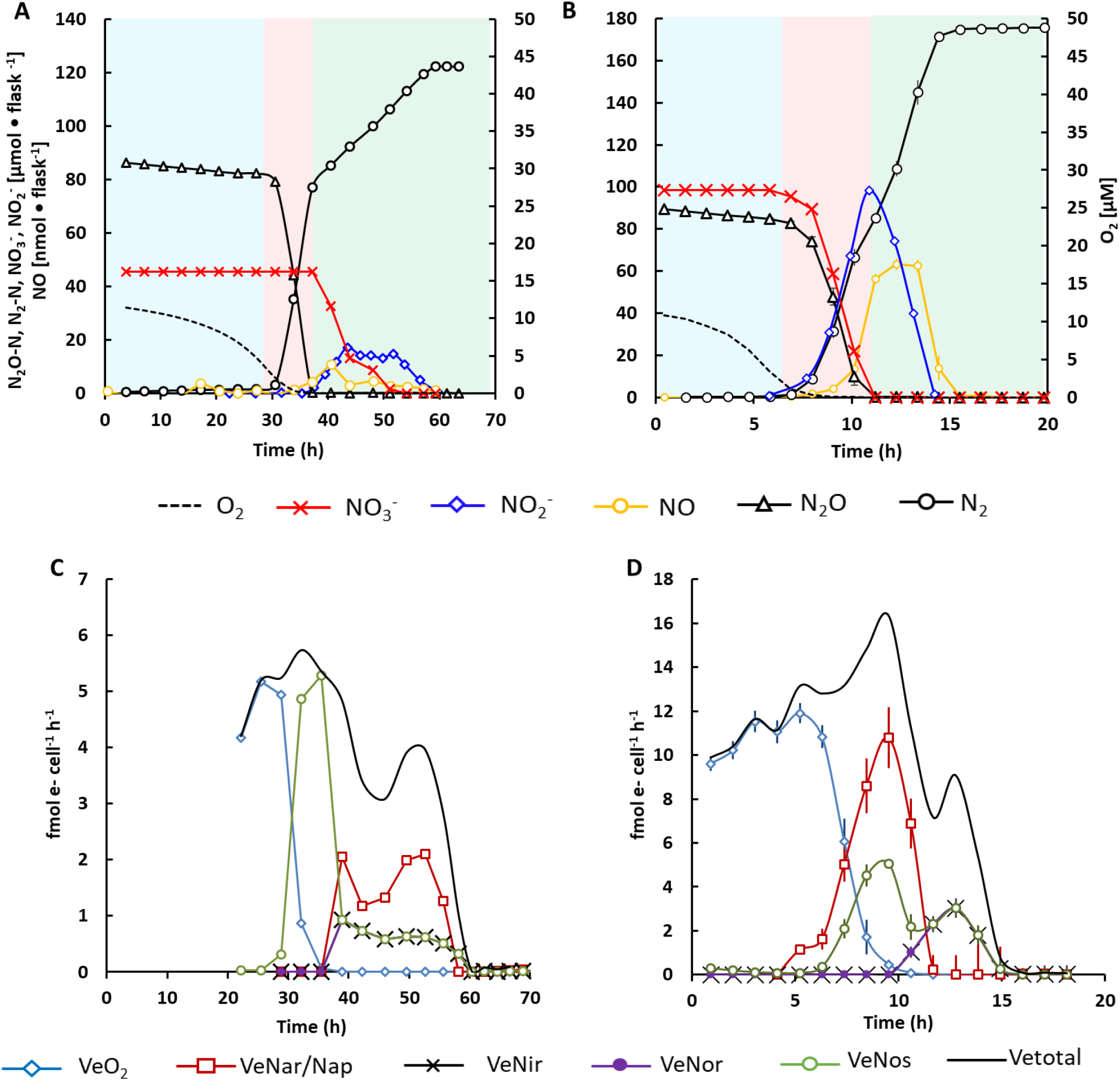
Competition for electrons between N_2_O- and NO_3_^−^ reductase; comparison of Nap- and Nar-carrying organisms. Gas kinetics and corresponding electron flow to oxidases (aerobic respiration) and to the different N-reductases (denitrification) in batch cultures of *Bradyrhizobium* strain HAMBI 2125 and *Paracoccus denitrificans*. Both organisms perform complete denitrification but *Bradyrhizobium* carries only the periplasmic NapA for dissimilatory NO_3_^−^ reduction while *P. denitrificans* carries both NapA and the membrane-bond NarG. The top panels (A, B) show the measured amounts of NO_2_^−^, NO, N_2_O and N_2_ per flask, and the concentration of O_2_ in the liquid. Note that the small but significant decrease in N_2_O during the aerobic phase was caused by sampling losses, not by N_2_O reduction or other reactions, as explained in Fig. S6. Background colors indicate the dominating respiratory activity. Blue: O_2_ respiration; Pink: N_2_O respiration; Green: Complete denitrification (NO_3_^−^ to N_2_). The bottom panels (C, D) show the calculated cell specific rates of electron flow (fmol cell^−1^ h^−1^) to the terminal oxidases (V_eO2_) and to the four reductases (V_eNar/Nap_, V_eNir_, V_eNor_ and V_eNos_). V_eNir_ and V_eNor_ were practically identical throughout (NO remained very low), thus indistinguishable in the panels. Strain HAMBI 2125 was incubated in 4 replicate flasks with YMB supplemented with 1 mM KNO_3_, of which only one is shown here (gas- and NO_2_^−^ kinetics of replicate flasks are shown in Fig. S3). *P. denitrificans* was incubated in 50 ml Sistrom’s medium with 2 mM KNO_3_ (n=7, bars show standard errors). At the start of the incubation, all flasks contained ~1% O_2_ in the headspace (~10 μM in liquid) and ~ 1.2 ml N_2_O (~90 μmol N_2_O-N). V_eNar/Nap_ and V_eNir_ represent rates of electron transport to NO_3_^−^ and NO_2_^−^, respectively (via NapA and NirK for *Bradyrhziobium* and via NarG+Nap and NirS for *P. denitrificans*).

The first phase, marked in blue, was dominated by O_2_ respiration. When the O_2_ concentration in the medium reached 4.6 μM, some NO_3_^−^ reduction took place, measured as a slight accumulation of NO_2_^−^ and NO (0.7 and 1.7 μmol; not clearly visible in the figure due to scaling). The small but significant decrease in N_2_O during the aerobic phase was caused by sampling losses, not by N_2_O reduction or other reactions, as explained in Fig. S6. The same is seen for experiments presented in Figs. 2B and S3-S5. Reduction of exogenous N_2_O started when the O_2_ concentration in the liquid was approximately 3.0 μM and rapidly reached a rate of 10.4 ± 1.2 μmol N flask^−1^ h^−1^, matching the corresponding N_2_ production rate of 9.7 ± 1.0 μmol N flask^−1^ h^−1^. This phase of rapid and stoichiometric reduction of N_2_O to N_2_, marked in pink, lasted until most of the exogenous N_2_O was reduced. No NO_3_^−^-reduction took place during reduction of the exogenous N_2_O. When the exogenous N_2_O was depleted, the rate of NO_3_-reduction increased and the rate of N_2_ production decreased instantaneously to 2.22 ± 0.18 μmol flask^−1^ h^−1^ (phase marked in green). When NO_3_^−^ reduction started, the NO_3_^−^ concentration increased sharply and reached approximately 200-350 μM (9.8-17.4 μmol flask^−1^), accounting for 20-35% of the initial 50 μmol NO_3_^−^-N. The N_2_ curve for strain HAMBI 2125 reached a stable plateau at 123 μmol N_2_-N, which accounts for the sum of N_2_O- and NO_3_-N present initially. The same experiment was conducted with other strains, but without measuring NO_2_^−^ (Figs S4, S5), and they all showed the same fast stoichiometric conversion of N_2_O to N_2_ and a subsequent slower production of N_2_ from NO_3_^−^, lasting until the N_2_O+NO_3_^−^-N was completely recovered as N_2_-N. Strain HAMBI 2125 controlled NO during the entire incubation period, reaching a small peak of 11 nmol flask^−1^ (6.9 nM in the liquid) around 41 hpi. For the other strains NO concentrations never exceeded 50 nM, and most strains kept NO concentrations <20 nM during the entire incubation (Figs 2A, S3, S4, S5).

### Nar carrying Paracoccus denitrificans showed no preference for N_2_O over NO_3_^−^

In accordance with our hypothesis, the Nar carrying *P. denitrificans* reduced NO_3_^−^ simultaneously with the exogenously supplied N_2_O, indicating no hampering of NO_3_^−^ reduction by N_2_O (Fig. 2B). In contrast to all the examined *Bradyrhizobium* strains (Figs 2A, S3, S4, S5), *P. denitrificans* started to reduce NO_3_^−^ and headspace N_2_O concomitantly when the O_2_ concentration was close to depletion and stoichiometrically accumulated NO_2_^−^ from NO_3_^−^ reduction. Nitrite reduction and emergence of NO started when most of the NO_3_^−^ had been reduced to NO_2_^−^.

### Cell specific electron flow rates

The measured gas- and NO_2_ kinetics were used to estimate the electron flow rates to the various reductase (μmol e^−^ flask^−1^ h^−1^) for each time increment between gas samplings. To convert these rates to cell specific electron flow rates (fmol cell^−1^ h^−1^), we needed estimates of the cell density throughout an incubation. For this, we used cumulated electron flow to the various electron acceptors (calculated from the gas- and NO_2_^−^-measurements), and the cell yield per mol electrons to each electron acceptor, which was determined in a separate experiment (see Experimental procedures and Fig. 5). In *Bradyrhizobium* strain HAMBI 2125 (Fig. 2C), the cell specific electron flow was sustained at a more or less constant level (5-6 fmol e^−^ cell^−1^ h^−1^) through the transition from respiring O_2_ to respiring N_2_O, until the external N_2_O was depleted (phases marked blue and pink in the corresponding gas kinetics Fig. 2A), but declined to ~3-4 fmol e^−^ cell^−1^ h^−1^ when depending on the activity of Nap. In *P. denitrificans* (Fig. 2D) there was instead a simultaneous electron delivery to Nar and Nos during the phase when exogenous N_2_O was being reduced (marked pink in the corresponding gas kinetics, Fig. 2B). This was later followed by electron flow to Nir, Nor and Nos, after the depletion of nitrate. The cell specific respiratory metabolism was 10-13 fmol e^−^ cell^−1^ h^−1^ during the phase dominated by aerobic respiration, then increased to 15-16 fmol e^−^ cell^−1^ h^−1^ when respiring NO_3_^−^ and N_2_O, followed by a decrease to approximately 8 fmol e^−^ cell^−1^ h^−1^ during respiration of the accumulated NO_2_-.

### Injection of N_2_O to actively denitrifying cultures resulted in an immediate arrest of NO_3_^−^reduction

To investigate whether the preferred electron flow to N_2_O in the foregoing experiments could be explained by regulation at the transcriptional level (i.e. early expression of *nos*), N_2_O was injected into a culture that was actively reducing NO_3_^−^ to N_2_. A transcriptional regulation would not result in an immediate switch to the exogenous N_2_O as electron acceptor but would instead be seen as a simultaneous reduction of all N-oxides present. This was tested on strain HAMBI 2125 (Fig. 3). The cultures were incubated in flasks containing YMB with 2 mM initial NO_3_^−^, and with 1% O_2_ but without exogenous N_2_O in the headspace. Denitrification (from NO_3_-reduction) commenced when O_2_ reached 4.3 μM and proceeded at a rate of 2.6 ± 0.4 μmol N flask^−1^ h^−1^. At 32 hpi, when the cells were actively reducing NO_3_^−^ to N_2_, approximately 300 μmol N_2_O-N was injected. This led to a complete suppression of NO_3_^−^ reduction and rapid decline in NO accumulation. The measured N_2_ production rate was similar to the measured N_2_O reduction rate (23.9 ±3.6 and 26.4 ±3.6 μmol N flask^−1^ h^−1^, respectively). The NO_3_^−^ reduction recovered upon depletion of external N_2_O, reflected in an increase in NO and an N_2_ production rate similar to that before the N_2_O injection. In an additional experiment, including all seven N_2_O-reducing strains, exogenous N_2_O was present in the headspace from the start of the incubation (Fig. S5). As expected from the first experiment (Fig. 2A), the exogenous N_2_O was rapidly reduced, followed by NO_3_^−^ reduction to N_2_ at a slower rate. A second injection of N_2_O into the actively denitrifying cultures resulted in an immediate increase in the N_2_ production rate, comparable with that during reduction of the initial N_2_O. These results supported the notion that N_2_O was preferred as electron acceptor over NO_3_^−^.

**Fig. 3.**
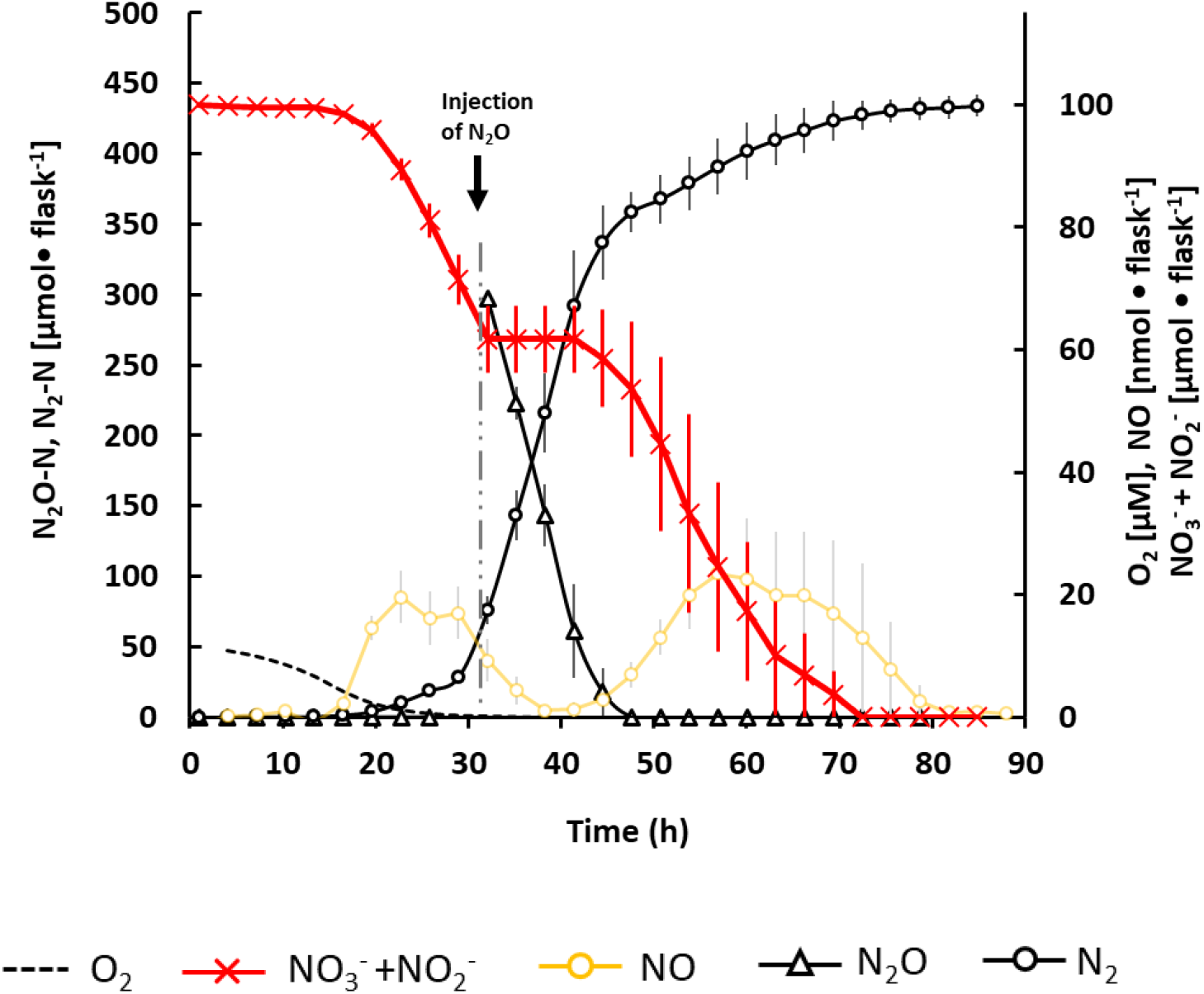
Effect of injecting N_2_O to an actively denitrifying culture. Gas kinetics for *Bradyrhizobium* strain HAMBI 2125 during and after transition from aerobic respiration to N-oxide respiration. Triplicate cultures were incubated in YMB medium supplemented with 2 mM KNO_3_ and with 0.7 ml O_2_ in the headspace (around 10 μM O_2_ in the medium). Approximately 4 ml N_2_O (~300 μmol N_2_O-N) were injected at 32 hpi, indicated by the arrow, into actively denitrifying cultures respiring NO_3_^−^. Bars show standard errors (n=3).

### The relative abundances of the reductases could not explain the preference for N_2_O over NO_3_^−^

A proteomics analysis was done for strain HAMBI 2125. The abundance of NapA, NirK and NosZ molecules, measured as intensities, were determined at different time points in cells sampled during aerobic respiration (21% O_2_), and during and after transition from 1% oxygen respiration to denitrification (Fig. 4). The level of NorC molecules detected was two orders of magnitude lower than those for the other reductases, suggesting that membrane-bound enzymes require a different extraction protocol. Therefore, only values for the periplasmic reductases NapA, NirK and NosZ were included here. The aerobic pre-incubation over several generations effectively removed most denitrification reductases that may have lingered from earlier periods of anoxia, seen from the signals from NapA and NirK being 293-585 times lower than those observed during denitrification. Nos values were only 16-47 times lower, indicating low levels of constitutive transcription of the *nos* gene and translation of the corresponding mRNA under aerobic conditions. All three reductases increased in abundance during the O_2_ depletion phase (blue) and when O_2_ approached depletion, the NapA/Nos and NapA/NirK ratios were 1.4±0.2 and 4.1±1.8 (sd, n=3). Interestingly, the NapA/Nos ratio was stable during the period of strong respiration of exogenous N_2_O, marked pink in Fig. 4, with values of 1.5± 0.3 and 1.2 ± 0.2, although practically no NO_3_^−^ reduction took place. The NapA/Nos ratio decreased to 0.7 during the phase of NO_3_^−^ reduction to N_2_ (after depletion of exogenous N_2_O). A similar pattern was found for the NapA/NirK ratios, while the NosZ/NirK ratios stayed around 1.0 during the anaerobic phase.

**Fig. 4.**
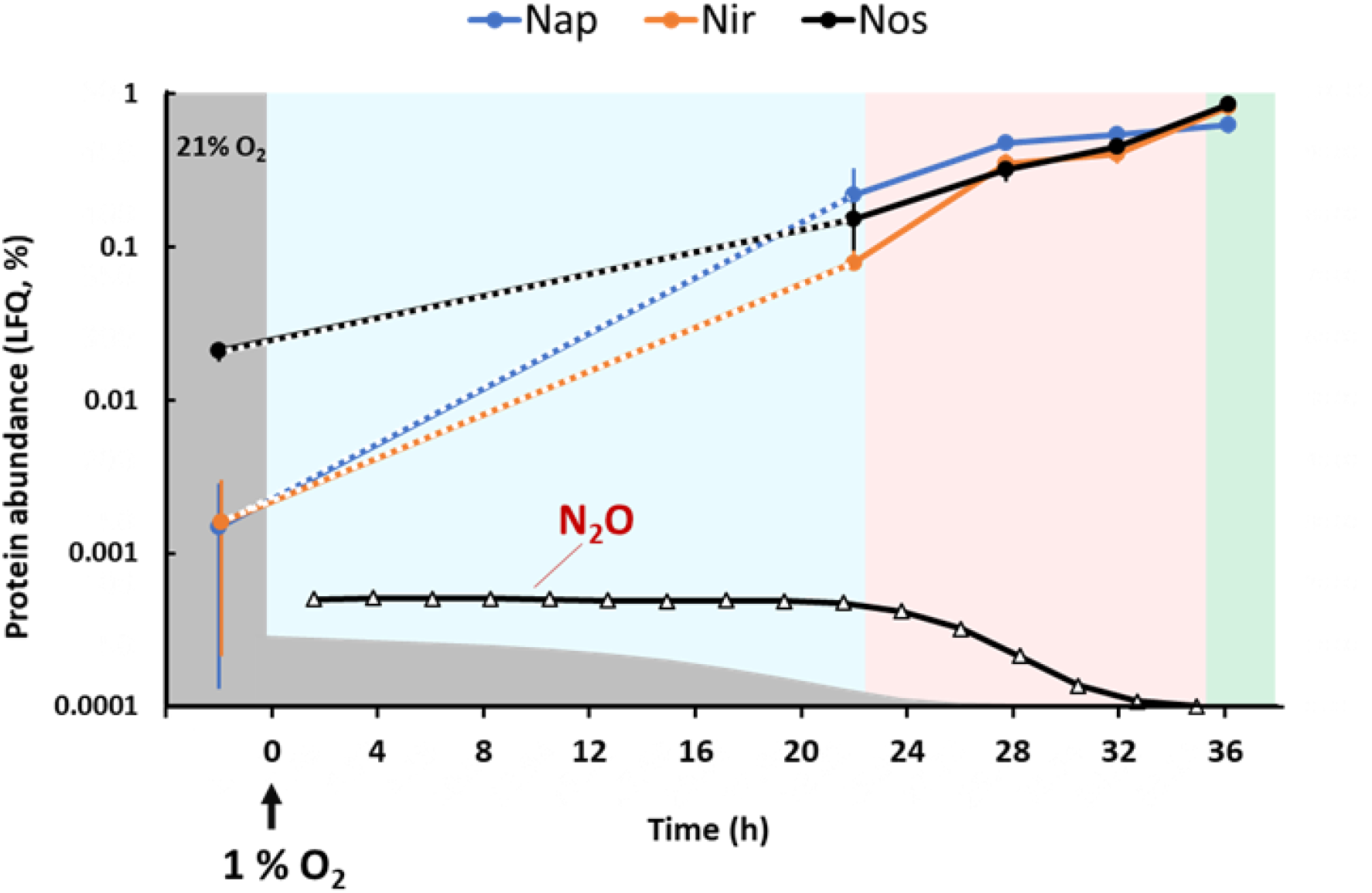
Abundance of denitrification reductases during and after transition from aerobic to anaerobic respiration. The abundances of Nap, NirK and Nos were determined by proteomics and calculated as percentages of the sum of all LFQ (label-free quantitation of proteins) abundances for each time point. The first samples were taken from triplicate flasks of *Bradyrhizobium* strain HAMBI 2125 growing under fully oxic conditions (21% O_2_). Subsamples from these precultures were inoculated into flasks with 1% O_2_, and samples for proteomics were taken when the cultures approached O_2_ depletion ([O_2_] = 5.8±0.2 μM) and during the anaerobic respiration. Concentrations of O_2_ (grey shading) and N_2_O (-Δ-) are schematically indicated here to illustrate the time of their depletion; for actual values of [O_2_] [N_2_O], [N_2_] and [NO_2_^−^], see Fig. S8. Background colors indicate the dominating respiratory activity. Blue: O_2_ respiration; Pink: N_2_O respiration; Green: Complete denitrification (NO_3_^−^ to N_2_). Bars show standard deviation (n=3).

### Growth rate and yield per mol electrons for aerobic and anaerobic respiration

The experiment was designed to determine the growth rate and growth yield per mol electrons for aerobic and anaerobic respiration (N_2_O and NO_3_^−^), but also to inspect if cumulated electron flow to the various electron acceptors could be used to predict cell density throughout a batch incubation. The results summarized in Fig. 5 confirm that cumulated electron flow can be used to predict increase in cell density (or dry weight, DW) throughout an incubation, provided that the yield per electron to the various electron acceptors is known. The aerobic growth rate was twice the anaerobic growth rate by N_2_O respiration (0.063 and 0.031 h^−1^). Growth from complete denitrification (NO_3_^−^ provided as e^−^ acceptor) was 0.024 h^−1^ (Fig. 5 A-C). The growth yield per mol electrons was 7.17±0.38 g DW mol^−1^ e^−^ for aerobic respiration, 5.1±0.10 g DW mol^−1^ e^−^ from N_2_O respiration and 4.03±0.20 g DW mol^−1^ e^−^ for complete denitrification (NO_3_^−^ to N_2_) (Fig. 5D). In the same experiment, we determined the DW per cell for cells grown by respiring O_2_, N_2_O and NO_3_^−^ (Fig. S7 B). Since the values for growth on the different electron acceptors were not significantly different, they were regarded as replicates, giving an average cell DW of 615 ±54 fg (sd, n=12).

**Fig. 5.**
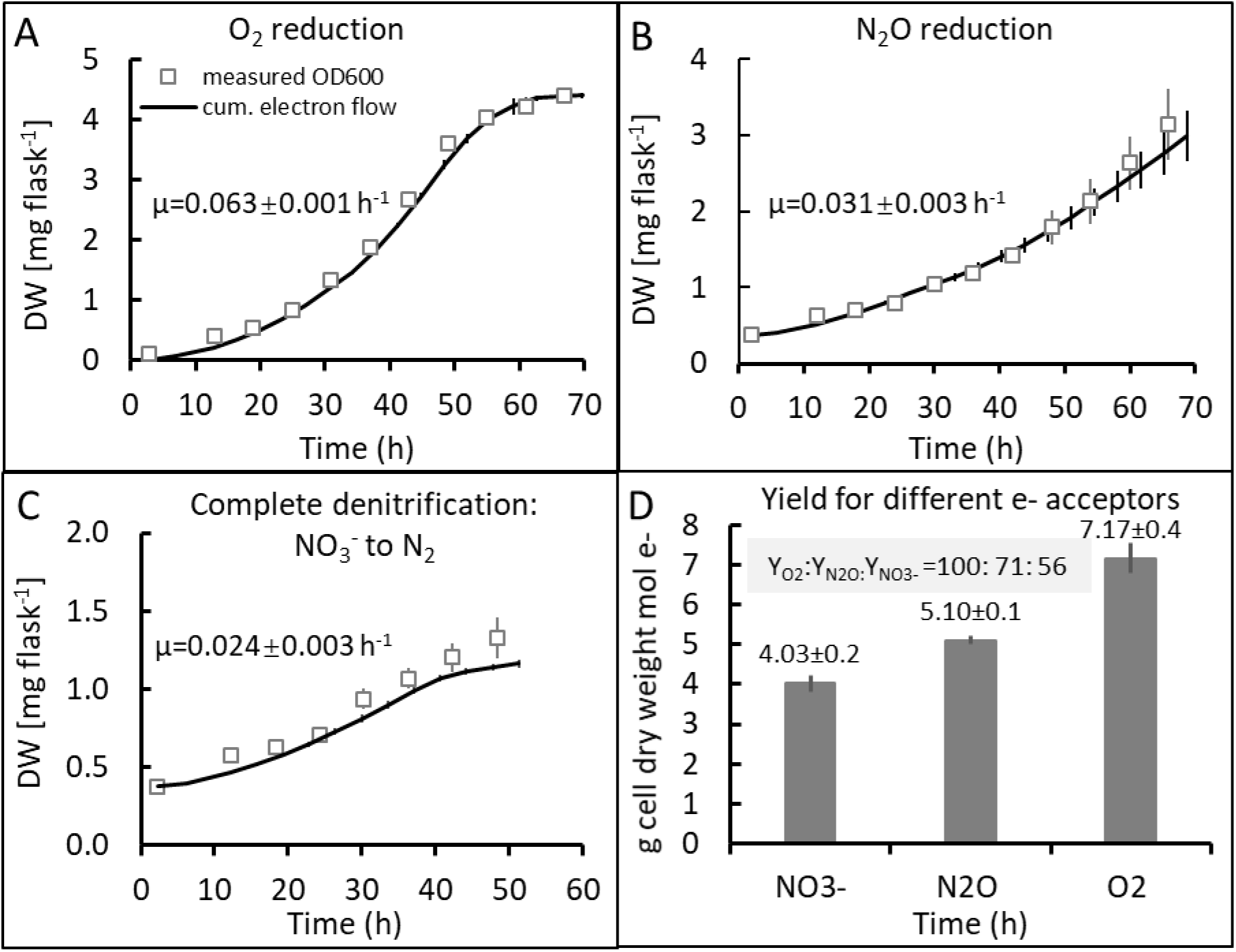
Growth and yield of *Bradyrhizobium* strain HAMBI 2125 when incubated in YMB at 28 °C and provided with O_2_, N_2_O or NO_3_^−^ as electron acceptor. All relevant gases were measured frequently using a robotized system. <u>Panels A-C</u> show estimated g dry weight (DW) flask^−1^ throughout the incubations either calculated from measured OD_600_ (□) using the factor 0.374 mg dry weight ml^−1^ OD_600_^−1^ (Fig. S7) or calculated from the measured cumulated electron flow (solid line) using measured yields per mol e^−^ in <u>panel D</u>. The specific growth rates (μ) were determined by regression of ln(OD_600_) against time during the period of exponential growth. <u>Panel A</u>. Growth using O_2_ as sole electron acceptor (initial amount 180 μmol flask^−1^). <u>Panel B</u>. Growth using N_2_O as sole electron acceptor (initial amount 800 μmol N flask^−1^). <u>Panel C</u>. Growth from complete denitrification (NO_3_^−^ to N_2_; initial amount of NO_3_^−^ was 200 μmol flask^−1^). <u>Panel D</u>. Yield per mol e^−^ obtained from growth using the different electron acceptors. The differences in yield were statistically different (p<0.01). O_2_ was depleted in the O_2_ treatment after 55 h, while N_2_O and NO_3_^−^ were not depleted during the incubations. Bars indicate standard deviation, n=3. Note: The initial value for DW, estimated by cumulated electron flow, was set equal to that based on measured OD.

## Discussion

The study comprised a set of taxonomically diverse *Bradyrhizobium* strains, as shown by the phylogenetic tree (Figs 1, S1). This genus is commonly found among the most dominant organisms in soil microbial communities (Delmont *et al*., 2012; Xu *at al*., 2014) and of large environmental and economic value for their N_2_-fixation ability in symbiosis with legumes. Based on the MLSA analysis of four concatenated housekeeping genes (Fig. 1) the strains clustered in eleven clades, covering various parts of the phylogenetic tree. Five strains in three clades did not group with described *Bradyrhizobium* species, suggesting that three new species may be delineated for these three clades. Our results from the denitrification analyses support other studies showing that complete denitrification pathways are more common in the *B. japonicum* supergroup than in other phylogenetic branches of *Bradyrhizobium* (Sameshima-Saito *et al.*, 2004; Mania *et al*., 2020). The complete denitrifiers in the present study belonged to clusters related to *B. shewense*, *B. ottawaense* and *B. guangxiense*, with the exception of one strain which clustered with the distantly related *B. lablabi* (Fig. 1). Although this suggests that some taxonomic groups of bradyrhizobia have higher frequency of complete denitrifiers than others, taxonomic position cannot be used as an indicator of the denitrification end-product since even very closely related strains may differ in this regard (Mania *et al.*, 2020).

Recent research has revealed several transcriptional and post-transcriptional mechanisms that influence the amounts of intermediate products released from denitrifying organisms (Liu *et al*., 2013; Torres *et al*., 2016; Lycus *et al*., 2018; Gaimster *et al*.,2019; Jiménez-Leiva *et al*., 2019). Although of value by itself, such information may be anecdotal. To understand the environmental relevance of a regulatory mechanism, it is important to know how widespread it is. We found the same preference for N_2_O over NO_3_^−^ as electron acceptor in all strains with complete denitrification, as did Mania *et al.* (2020) for a different set of isolates. This indicates a phenomenon common to bradyrhizobia. In theory, it could be explained by lower numbers of NapA molecules compared to NosZ molecules. The results by Mania *et al.* (2020) for strain AC 87j1, which is closely related to strain HAMBI 2125, could not disprove this, since transcription analysis showed 5-8 times lower copy numbers of transcripts coding for NapA compared to NosZ. This prompted us to perform a proteomics analysis in the present study. The results (Fig. 4) provided clear evidence that the strong N_2_O reduction was not due to higher abundance of NosZ *vs* NapA, thus supporting the hypothesis by Mania *et al.* (2020) that the electron pathway to NosZ is inherently more competitive than that to NapA.

Electrons from the quinol pool are channeled to Nap via the membrane-bound NapC, and to Nos, Nor and Nir via the membrane-bound *bc*1 complex and periplasmic cytochromes (Shapleigh, 2013). Most likely, the electron pathway to NosZ via *bc*1 is inherently more competitive than that to NapAB via NapC, plausibly due to a combination of stronger affinity for quinol and a higher k_cat_ of *bc*_1_ versus NapC. This implies a general mechanism which may apply not only to bradyrhizobia, but possibly to all denitrifiers that carry Nap and Nos but lack the membrane-bound Nar. Verifying this will require testing of a wide range of organisms. As a first step, we included in this study a comparison of the denitrification kinetics of the bradyrhizobia with *P. denitrificans*, which carries Nar in addition to Nap and Nos. The results supported our hypothesis by clearly demonstrating that, while the *Bradyrhizobium* strains left NO_3_^−^ untouched as long as exogenously supplied N_2_O was present (Figs 2A, S3, S4, S5), *P. denitrificans* reduced these electron acceptors simultaneously (Fig. 2B). This suggests that the electron pathways from the quinol pool to NarG via NarI and NarH and to NosZ via the *bc*_1_ complex and cytochrome c compete equally well for electrons, at least in *P. denitrificans*.

The proteomics analysis of HAMBI strain 2125 showed that NosZ was synthesized under fully aeriated conditions, while NapA and NirK were not. Before sampling for proteomics analysis, the cultures had been incubated in flasks with vigorous stirring for at least 10 generations to secure that denitrification reductases produced during earlier exposure to denitrifying conditions were diluted to near extinction in the cells. This was the case for NapA and NirK in the sample from 21% O_2_ (Fig. 4). NosZ molecules were, however, approximately 10 times more abundant than NapA and NirK during this phase, which suggests constitutive, albeit low, synthesis of NosZ. The transcription analysis of strain AC87j1 reported by Mania *et al.* (2020) provides no clear explanation to this, though. In the presence of 5 μM O_2_ in the liquid, there were < 0.3 *nosZ* and *nirK* transcripts cell^−1^ with only slightly higher levels of *nosZ* than *nirK*. The *napA* transcription was somewhat higher with 0.8 copies cell^−1^. It is possible, still, that a low but steady production of Nos takes place in these organisms irrespective of O_2_ concentrations, in line with the higher β-galactosidase activity observed for a *nos-lacZ* fusion of *B. japonicum* compared to a *nap-lacZ* fusion (Bueno *et al.*, 2017; Torres *et al.*, 2017). This could be a strategy for cells to be prepared for anaerobic conditions and avoid being entrapped in anoxia without enough energy to produce the denitrification machinery (Lycus *et al.*, 2018). If so, it suggests that these bacteria may be ready to reduce any N_2_O produced by other organisms in their environment, thus acting as net sinks for this greenhouse gas.

Seven strains lacked the last step of denitrification and thus had N_2_O as end-product, explained by the absence of the entire *nos* operon in all cases (Table S2). When exposed to denitrifying conditions in the environment, such organisms will reduce all available N-oxides to N_2_O and thus act as net producers of this greenhouse gas. Three strains had NO as denitrification end-product, but only HAMBI 2149 lacked the entire *nor* operon (Table S2). The other two strains, HAMBI 2153 and 2299, contained *nor* operons but showed some sequence differences compared to strains with functional NO reduction. Further investigations of the lacking NO reduction were, however, beyond the scope of the present study. Organisms that produce NO but are incapable of reducing it are apparently not uncommon (Casella *et al*., 2006; Falk *et al.*, 2010; Mania *et al.*, 2020). In the environment such organisms are most likely found in consortia with organisms that express Nor or other NO-reducing enzymes.

The estimated growth yields were calculated based on the electron flow to the different reductases (Fig. 5) and the proportions were expected to match that of the charge separations per electron transport for the various respiratory pathways, as listed by (van Spanning *et al.*, 2007). The number of charge separations per 10 electrons (q/10e^−^) are 50 for low affinity oxidase, 40 for high affinity oxidases, 30 for Nos alone, and 26 for the entire Nap, Nir, Nor, Nos denitrification pathway (NO_3_^−^ to N_2_), which would a yield ratio of 100:60:52 for the three pathways (O_2_, N_2_O, NO_3_→N_2_) if we assume that low affinity oxidases are used, and 100:75:65 for high affinity oxidases. Our measured yields were 12, 8.5 and 6.7 *10^12^ cells per mol electrons for the three pathways, i.e. a yield ratio of 100:71:56, which agrees reasonably with that predicted for high affinity oxidases.

The emerging evidence that bradyrhizobia with complete denitrification deplete N_2_O in their surroundings before reducing available NO_3_^−^ is promising from an environmental point of view since they have the potential to act as net sinks for N_2_O produced by other organisms in their vicinity. In legume fields, nodules and other N-rich organic material will be degraded, releasing NH_4_^+^, which will rapidly be metabolized by nitrifying bacteria and archaea, producing NO_3_^−^ while consuming O_2_. This will create optimal conditions for denitrification. Legume cropped fields account for a significant share of the global N_2_O emissions (Jensen *et al.*, 2012), and it is likely that coupled nitrification-denitrification is a main cause (Bakken and Frostegård, 2017). One way of mitigating these emissions is to enhance the number of efficient N_2_O reducing bacteria in the soil microbial community. Rhizobial inoculants are used since long to increase the yield of leguminous crops and at the same time avoid the need for synthetic fertilizers. Different combinations of rhizobial strain and legume cultivar may differ substantially in nitrogen fixation effectiveness, which is also affected by environmental conditions (Woliy *et al.*, 2019; Lindström and Mousavi, 2020), and large efforts are put into the development of new inoculants that are compatible with various types of legumes as well as soil types and climates (Martínez *et al.*, 2016). The present study shows that *Bradyrhizobium* strains also differ with respect to N_2_O reduction, and suggests that new inoculants should be selected among rhizobia that express Nos.

## Experimental Procedures

### Bacterial strains

The rhizobial isolates were obtained from various culture collections. Table 1 shows a list of host plant and geographic origin of the strains used in the phylogenetic analysis (all listed isolates; details see (Van Rossum *et al*., 1993; Van Rossum *et al*., 1994; Zhang *et al*., 1999; Chang *et al*., 2011; Wang *et al*.,2013; Li *et al*., 2015; Degefu *et al*., 2017) and the strains analyzed for their denitrification capacity (all except the four strains from Ethiopia earlier determined to have a complete denitrification pathway (Mania *et al.* 2020). Preparation of *Bradyrhizobium* cultures and growth conditions followed Mania *et al.* (2020). All cultures were grown at 28 °C in Yeast Mannitol Broth (YMB; 10 g l^−1^ D-mannitol, 0.5 g l^−1^ K_2_HPO_4_, 0.2 g l^−1^ MgSO_4_·7H_2_O, 0.1 g l^−1^ NaCl and 0.5 g l^−1^ yeast extract, pH 7.0; Schwartz, 1972).

### Phylogenetic analyses

DNA was extracted using QIAamp DNA Mini Kit (Qiagen). The genome sequences were obtained from another ongoing project: (JGI Gold study ID: Gs0134353), and their Genome accession numbers are listed in Table S3. Genome sequences were analyzed using RAST (Rapid Annotations using Subsystems Technology). Whole-genome average nucleotide identity (ANI) comparisons were done using FastANI v.0.1.2. A phylogenetic tree that included all HAMBI and CCBAU strains from this study (Table 1), as well as the Ethiopian strains AC70c, AC86d2, AC87j1 and AC101b and the type strain of all species of the genus *Bradyrhizobium* (February 2020, http://www.bacterio.net/bradyrhizobium.html), was constructed based on multilocus sequence analysis (MLSA) of the four housekeeping genes *atpD* (ATP synthase F1, beta subunit), *glnII* (glutamine synthetase type II), *recA* (recombinase A), *rpoB* (RNA polymerase, beta subunit). The sequences from all HAMBI strains used in the MLSA analysis were obtained from the present study. Amplicons of *rpoB* and *atpD* of the Ethiopian strains AC70c, AC86d2 and AC101b and *recA* from AC70c were Sanger sequenced. For primers and PCR conditions, see Table S4. All other sequences were obtained from GenBank (http://www.ncbi.nlm.nih.gov/genbank). The GeneBank accession numbers for all housekeeping genes used are listed in Table S5. The gene sequences were aligned by ClustalW (Larkin *et al.*, 2007) in Bioedit v7.0.5.3 (Hall, 1999), and the alignments were edited manually. The best-fit models of nucleotide substitution were selected based on Akaike information criterion (AIC) applying MEGA7 (Kumar *et al.*, 2016). The Bayesian phylogenetic analyses of the concatenated loci (*atpD*-*recA*-*glnII*-*rpoB*) of 78 *Bradyrhizobium* strains were performed using the algorithm Metropolis coupled Markov chain Monte Carlo (MCMCMC) twice for ten million generations using MrBayes v3.2.7a program (Ronquist *et al.*, 2012). *Neorhizobium galegae* was used to root the tree. The general time reversible plus gamma distribution plus invariable sites (GTR + G + I) was chosen as the best-fit model of phylogenetic analyses of the four partition of the concatenated dataset (*atpD-glnII-recA-rpoB*). The algorithms were analyzed using Tracer v1.7 (Rambaut *et al.*, 2018), and the phylogenetic trees were visualized by Figtree v1.4.3 (https://beast.community/figtree). For comparison with the MLSA tree based on four housekeeping genes, a phylogenetic tree of all the HAMBI test strains listed in Table S2 was constructed using the Genome Taxonomy Database Toolkit (GTDB-Tk) v0.3.2, which is based on a phylogeny inferred from the concatenation of 120 ubiquitous, single-copy proteins (Chaumeil *et al*., 2020).

### Denitrification end products and gas kinetics analyses

Prior to these experiments the cultures were streaked for purity and single colonies were picked and checked for possible contaminating bacteria by sequencing of the 16S rRNA genes (for primers and PCR conditions see Table S4). Pure cultures were preserved as in glycerol (15%) at −80 °C. Cultures were raised from these frozen stocks in stirred batches of 50 ml YMB supplemented with filter sterilized KNO_3_ and/or KNO_2_ solutions. The flasks were sealed with sterilized, gas tight butyl rubber septa (Matrix AS, Norway). Air was removed by applying vacuum for 6 x 360 s, followed by 30 s He-filling. The desired amount of O_2_ and/or N_2_O was injected into the headspace, and the over-pressure released (Molstad *et al.*, 2007). Denitrification kinetics of *Bradyrhizobium* strains with complete denitrification pathways were compared to the model bacterium *Paracoccus denitrificans* PD1222, which was grown under the same conditions as the *Bradyrhizobium* strains, except that Sistrom’s medium supplemented with 2 mM KNO_3_ was used (Sistrom, 1962; Bergaust *et al.*, 2010).

Denitrification end-products were determined for all strains as described previously (Lycus *et al.*, 2017). Detailed gas kinetics were measured following Mania *et al*. (2020). Pre-cultures were added to incubation flasks (n=3) supplied with 1 mM initial KNO_3_ and, when indicated, also with 0.5 mM KNO_2_. Unless stated otherwise, 0.7 mL O_2_ (1 vol% in headspace) and 0.7 ml N_2_O (approximately 50 μmol N flask^−1^) were added to the headspace at the start of the experiment (0 hpi, hours post inoculation). For the determination of end-products, the amounts of O_2_, NO, N_2_O and N_2_ in the headspace were measured at the start and after 10 days of incubation.

All strains with a complete denitrification pathway according to the end-point analysis, as well as all HAMBI strains that had NO as end-product, were investigated further by analyzing gas kinetics in response to oxygen depletion. To secure an inoculum in which any previously synthesized denitrification enzymes had been diluted out by aerobic growth, all cultures were precultured aerobically for at least 6-8 generations. This was done in uncapped flasks covered with aluminum foil (to allow oxygen transport), with vigorous stirring (600 rpm) at 28 ^°^C. To further ensure the absence of hypoxia, the cultures were transferred to new flasks before the OD_600_ reached 0.3. Portions of the precultures were inoculated into 120 ml flasks containing 50 ml YMB supplemented with 1, 2 or 3 mM KNO_3_ and with He, O_2_ and N_2_O in headspace as described above for the end-product measurements. The flasks were placed in a robotized incubation system that frequently measured O_2_, NO, N_2_O and N_2_ in headspace (Molstad *et al.*, 2007; Molstad *et al.*, 2016). Leakage and sampling losses were taken into account when calculating the rates of production/consumption of each gas, see Molstad *et al*. (2007) for details. Earlier experiments determined that no gas production (NO, N_2_O or N_2_) took place from chemical reactions between of NO_2_^−^ and the medium under the conditions used (Mania *et al.*, 2020).

### Nitrite measurements

Samples for NO_2_^−^ measurements (0.1-1.0 ml) were taken manually from the liquid phase through the rubber septum of the flasks using a sterile syringe. NO_2_^−^ concentrations were determined immediately after each sampling by drawing a 10 μL liquid sample which was injected into a purging device containing 50% acetic acid with 1% (w/v) NaI (at room temperature), by which NO_2_^−^ is reduced instantaneously to NO (Cox, 1980; MacArthur *et al.*, 2007). The quantities of resultant NO were measured by chemiluminescence using a NO analyser (Sievers NOA™ 280i, GE Analytical Instruments). The system was calibrated by injecting standards. Sampling for NO_2_^−^ measurements was done as frequently as the gas sampling.

### Proteomics

Relative quantification of the denitrification reductases was done for the strain *Bradyrhizobium* sp. HAMBI 2125, chosen as a representative for the complete denitrifiers (Fig. 1), which all showed the same preference for N_2_O over NO_3_^−^ reduction (Figs 2A, S3, S4, S5). Cells were first raised from frozen stock under strictly aerobic conditions as explained above. As the aerobic preculture reached OD_600_ = 0.05 (2.9*10^7^ cells ml^−1^), it was used to inoculate 18 flasks (1 ml inoculum flask^−1^) containing YMB with 1 mM NO_3_^−^ and with 1 ml N_2_O (approximately 80 μmol N_2_O-N flask^−1^) and 1 % O_2_ in headspace. Samples for protein analysis were taken i) from the aerobic inoculum; ii) just before transition to anaerobic respiration (22 hpi, when O_2_ reached ~4.6 μM); iii) during the period of rapid reduction of exogenous N_2_O (27 and 32 hpi); and iv) during the period of NO_3_^−^ reduction to N_2_ (36 hpi). Three replicate flasks were harvested at each time point. Gas kinetics, NO_2_^−^ concentrations and sampling times for proteomics are shown in Fig. S8. After measuring the OD_600_, the samples were centrifuged (10 000 g, 4 °C, 10 min) and the cell pellets were stored at −20 °C for protein extraction. In addition, we harvested and froze cells from the aerobic preculture.

To extract the proteins, the cell pellets were thawed, resuspended in lysis buffer (20 mM Tris-HCl pH8, 0.1 % v/v Triton X-100, 200 mM NaCl, 1 mM DTT) and treated with 3×45 s bead beating with glass beads (particle size≤106 μm Sigma) at maximum power and cooling on ice between the cycles (MP Biomedicals™ FastPrep-24™, Thermo Fischer Scientific Inc). Cell debris was removed by centrifugation (10 000 g; 5 min) and the supernatant, containing water soluble proteins, was used for proteomic analysis based on Orbitrap-MS as described before (Conthe *et al.*, 2019). To quantify the denitrification reductases we calculated the fraction of the individual enzymes as percentages of the sum of all LFQ (label-free quantitation of proteins) abundances for each time point.

### Determinations of growth and yield

Aerobic and anaerobic growth was investigated in detail for *Bradyrhizobium* sp. HAMBI 2125 to determine the growth yield per mol electrons transferred to O_2_, N_2_O, and to the entire denitrification pathway. Four treatments were compared: 1) 7% O_2_ in headspace; 2) 0% O_2_, 4 mM NO_3_^−^; 3) 0% O_2_ and 10 ml N_2_O; 4) 7% O_2_, 4 mM NO_3_^−^ and 10 ml N_2_O. All treatments had an excess of electron donor (>70 times more electrons than needed to reduce the electron acceptor). Pre-cultures were started from single colonies and grown in YMB under fully oxic conditions (21% O_2_) for inoculation of the aerobic batch cultures (treatments 1 and 4), and under anoxic conditions for inoculation of the anaerobic batches (treatments 2 and 3). The OD_600_ of the aerobic pre-cultures was kept < 0.3 by regularly inoculating portions of the culture into flasks with fresh medium. There were 6 replicate flasks for each treatment. Liquid samples were taken from three of them every 6 h (0.85 ml) for growth measurements (OD_600_). The three other replicates were only monitored for gas kinetics and were then used to measure final cell dry weight, to assess the growth yield per mol electrons to O_2_ and N-oxides. For determinination of cell dry weight, cultures were centrifuged, washed twice in distilled water and dried at 100 ^o^C before weighing. Cell numbers were determined by microscopic counts at selected sampling times, which gave a conversion factor of 5.8*10^8^ cells ml^−1^ * OD_600_^−1^. The estimated yield from growth on O_2_, N_2_O and NO_3_^−^ was 12.0, 8.5 and 6.7 E12 cells mol^−1^ electrons, respectively (Figs 5, S7). The yields were used to calculate cell density throughout the incubations, enabling the calculation of cell specific rates of electron flows to the various electron acceptors, for details see Fig. S7.

## Supporting information

Supplementary Figures and Tables_Gao et al

## Acknowledgements

This project was supported by Kingenta Ecological Engineering Co., LTD. Yuan Gao is grateful to the China Scholarship Council (CSC) for financial support. We thank Dr. Xinghua Sui from China Agricultural University for providing seven of the *Bradyrhziobium* strains.

## Ethics declarations

The authors declare that they have no conflict of interest.

## References

Akiyama, H., Hoshino, Y.T., Itakura, M., Shimomura, Y., Wang, Y., Yamamoto, A., et al. (2016) Mitigation of soil N_2_O emission by inoculation with a mixed culture of indigenous *Bradyrhizobium diazoefficiens*. Sci Rep 6: 32869.

Arya, S.S., Salve, A.R., Chauhan, S. (2016) Peanuts as functional food: a review. J Food Sci Technol 53: 31–41.

Avontuur, J.R., Palmer, M., Beukes, C.W., Chan, W.Y., Coetzee, M.P.A., Blom, J., et al. (2019) Genome-informed *Bradyrhizobium* taxonomy: where to from here? Syst Appl Microbiol 42: 427–439.

Bakken, L.R., Frostegård, Å. (2017) Sources and sinks for N_2_O, can microbiologist help to mitigate N_2_O emissions? Environ Microbiol 19: 4801–4805.

Bedmar, E.J., Robles, E.F., Delgado, M.J. (2005) The complete denitrification pathway of the symbiotic, nitrogen-fixing bacterium *Bradyrhizobium japonicum*. Biochem Soc Trans 33: 141–144.

Bergaust, L., Mao, Y., Bakken, L.R., Frostegård, Å. (2010) Denitrification response patterns during the transition to anoxic respiration and posttranscriptional effects of suboptimal ph on nitrogen oxide reductase in *Paracoccus denitrificans*. Appl Environ Microbiol 76: 6387–6396.

Berks, B.C., Ferguson, S.J., Moir, J.W.B., Richardson, D.J. (1995) Enzymes and associated electron transport systems that catalyse the respiratory reduction of nitrogen oxides and oxyanions. Biochim Biophys Acta 1232: 97–173.

Bueno, E., Robles, E.F., Torres, M.J., Krell, T., Bedmar, E.J., Delgado, M.J., et al. (2017) Disparate response to microoxia and nitrogen oxides of the *Bradyrhizobium japonicum napEDABC*, *nirK* and *norCBQD* denitrification genes. Nitric Oxide 68: 137–149.

Casella, S., Shapleigh, J.P., Toffanin, A., Basaglia, M. (2006) Investigation into the role of the truncated denitrification chain in *Rhizobium sullae* strain HCNT1. Biochem Soc Trans 34: 130–132.

Chang, Y.L., Wang, E.T., Sui, X.H., Zhang, X.X., Chen, W.X. (2011) Molecular diversity and phylogeny of rhizobia associated with *Lablab purpureus* (Linn.) grown in Southen China. Syst Appl Microbiol 34: 276–284.

Chaumeil, P.A., Mussig, A.J., Hugenholtz, P., Parks, D.H. (2020) GTDB-Tk: a toolkit to classify genomes with the Genome Taxonomy Database. Bioinformatics 36: 1925–1927.

Conthe, M., Lycus, P., Arntzen, M.Ø., Ramos da Silva, A., Frostegård, Å., Bakken, L.R., et al. (2019) Denitrification as an N_2_O sink. Water Res 151: 381–387.

Cox, R.D. (1980) Determination of nitrate and nitrite at the parts per billion level by chemiluminescence. Anal Chem 52: 332–335.

Davidson, E.A., Kanter, D. (2014) Inventories and scenarios of nitrous oxide emissions. Environ Res Lett 9: 105012.

Degefu, T., Wolde-meskel, E., Woliy, K., Frostegård, Å. (2017) Phylogenetically diverse groups of *Bradyrhizobium* isolated from nodules of tree and annual legume species growing in Ethiopia. Syst Appl Microbiol 40: 205–214.

Delgado, M.J., Bonnard, N., Tresierra-Ayala, A., Bedmar, E.J., Müller, P. (2003) The *Bradyrhizobium japonicum napEDABC* genes encoding the periplasmic nitrate reductase are essential for nitrate respiration. Microbiology 149: 3395–3403.

Delmont, T.O., Prestat, E., Keegan, K.P., Faubladier, M., Robe, P., Clark, I.M., et al. (2012) Structure, fluctuation and magnitude of a natural grassland soil metagenome. ISME J 6: 1677–1687.

Falk, S., Liu, B., Braker, G. (2010) Isolation, genetic and functional characterization of novel soil *nirK*-type denitrifiers. Syst Appl Microbiol 33: 337–347.

Gaimster, H., Hews, C.L., Griffiths, R., Soriano-Laguna, M.J., Alston, M., Richardson, D.J., et al. (2019) A central small RNA regulatory circuit controlling bacterial denitrification and N_2_O emissions. mBio 10: e01165–19.

Hall, T.A. (1999) BioEdit: a user-friendly biological sequence alignment editor and analysis program for Windows 95/98/NT. Nucleic Acids Symposium Series 41: 95–98.

Hénault, C., Revellin, C. (2011) Inoculants of leguminous crops for mitigating soil emissions of the greenhouse gas nitrous oxide. Plant Soil 346: 289–296.

Itakura, M., Uchida, Y., Akiyama, H., Hoshino, Y.T., Shimomura, Y., Morimoto, S., et al. (2013) Mitigation of nitrous oxide emissions from soils by *Bradyrhizobium japonicum* inoculation. Nat Clim Chang 3: 208–212.

Jensen, E.S., Peoples, M.B., Boddey, R.M., Gresshoff, P.M., Hauggaard-Nielsen, H., Alves, J.R., et al. (2012) Legumes for mitigation of climate change and the provision of feedstock for biofuels and biorefineries. A review. Agron Sustain Dev. 32: 329–364.

Jiménez-Leiva, A., Cabrera, J.J., Bueno, E., Torres, M.J., Salazar, S., Bedmar, E.J., et al. (2019) Expanding the Regulon of the *Bradyrhizobium diazoefficiens* NnrR Transcription Factor: New Insights Into the Denitrification Pathway. Front Microbiol 10: 1926.

Kumar, S., Stecher, G., Tamura, K. (2016) MEGA7: Molecular Evolutionary Genetics Analysis Version 7.0 for Bigger Datasets. Mol Biol Evol 33: 1870–1874.

Larkin, M.A., Blackshields, G., Brown, N.P., Chenna, R., Mcgettigan, P.A., McWilliam, H., et al. (2007) Clustal W and Clustal X version 2.0. Bioinformatics 23: 2947–2948.

Li, Y.H., Wang, R., Zhang, X.X., Young, P.W., Wang, E.T., Sui, X.H., et al. (2015) *Bradyrhizobium guangdongense* sp. nov. and *Bradyrhizobium guangxiense* sp.nov., isolated from effectvie nodules of peanut. Int J Syst Evol Microbiol 65: 4655–4661.

Lindström, K., Mousavi, S.A. (2020) Effectiveness of nitrogen fixation in rhizobia. Microb Biotechnol 13: 1314–1335.

Liu, B., Mao, Y., Bergaust, L., Bakken, L.R., Frostegård, Å. (2013) Strains in the genus *Thauera* exhibit remarkably different denitrification regulatory phenotypes. Environ Microbiol 15: 2816–2828.

Lycus, P., Bøthun, K.L., Bergaust, L., Shapleigh, J.P., Bakken, L.R., Frostegård, Å. (2017) Phenotypic and genotypic richness of denitrifiers revealed by a novel isolation strategy. ISME J 11: 2219–2232.

Lycus, P., Soriano-Laguna, M.J., Kjos, M., Richardson, D.J., Gates, A.J., Milligan, D.A., et al. (2018) A bet-hedging strategy for denitrifying bacteria curtails their release of N_2_O. Proc Natl Acad Sci 115: 11820–11825.

MacArthur, P.H., Shiva, S., Gladwin, M.T. (2007) Measurement of circulating nitrite and S-nitrosothiols by reductive chemiluminescence. J Chromatogr B 851: 93–105.

Mania, D., Woliy, K., Degefu, T., Frostegård, Å. (2020) A common mechanism for efficient N_2_O reduction in diverse isolates of nodule-forming bradyrhizobia. Environ Microbiol 22: 17–31.

Martínez, J., Negrete-Yankelevich, S., Godinez, L.G., Reyes, J., Esposti, M.D., Martínez Romero, E. (2016) Short-Term Evolution of Rhizobial Strains Toward Sustainability in Agriculture. In Microbial Models: From Environmental to Industrial Sustainability. Springer: Singapore, pp. 277–292.

Mesa, S., Velasco, L., Manzanera, M.E., Delgado, M.J., Bedmar, E.J. (2002) Characterization of the *norCBQD* genes, encoding nitric oxide reductase, in the nitrogen fixing bacterium *Bradyrhizobium japonicum*. Microbiology 148: 3553–3560.

Molstad, L., Dörsch, P., Bakken, L.R. (2016) Improved robotized incubation system for gas kinetics in batch cultures. ResearchGate. https://doi.org/10.13140/RG.2.2.30688.07680.

Molstad, L., Dörsch, P., Bakken, L.R. (2007) Robotized incubation system for monitoring gases (O_2_, NO, N_2_O N_2_) in denitrifying cultures. J Microbiol Methods 71: 202–211.

Obando, M., Correa-Galeote, D., Castellano-Hinojosa, A., Gualpa, J., Hidalgo, A., Alché, J.D., et al. (2019) Analysis of the denitrification pathway and greenhouse gases emissions in *Bradyrhizobium* sp. strains used as biofertilizers in South America. J Appl Microbiol 127: 739–749.

Rambaut, A., Drummond, A.J., Xie, D., Baele, G., Suchard, M.A. (2018) Posterior summarization in Bayesian phylogenetics using Tracer 1.7. Syst Biol 67: 901–904.

Reay, D.S., Davidson, E.A., Smith, K.A., Smith, P., Melillo, J.M., Dentener, F., et al. (2012) Global agriculture and nitrous oxide emissions. Nat Clim Chang 2: 410–416.

Richardson, D.J. (2000) Bacterial respiration: a flexible process for a changing environment. Microbiology 146: 551–571.

Richardson, D.J., Berks, B.C., Russell, D.A., Spiro, S., Taylor, C.J. (2001) Functional, biochemical and genetic diversity of prokaryotic nitrate reductases. Cell Mol Life Sci 58: 165–178.

Rogelj, J., Shindell, D., Jiang, K., Fifita, S., Forster, P., Ginzburg, V., et al. (2018) Mitigation pathways compatible with 1.5 °C in the context of sustainable development. In, Masson-Delmotte V, Zhai P, Pörtner HO, Roberts D, Skea J, Shukla PR, et al. (eds). Global Warming of 1.5 °C. An IPCC Special Report on the impacts of global warming of 1.5°C above pre-industrial levels and related global greenhouse gas emission pathways, in the context of strengthening the global response to the threat ofclimate change, sustainable development, and efforts to eradicate poverty; https://www.ipcc.ch/site/assets/uploads/sites/2/2019/02/SR15_Chapter2_Low_Res.pdf.

Ronquist, F., Teslenko, M., Van Der Mark, P., Ayres, D.L., Darling, A., Höhna, S., et al. (2012) Mrbayes 3.2: Efficient bayesian phylogenetic inference and model choice across a large model space. Syst Biol 61: 539–542.

Sameshima-Saito, R., Chiba, K., Minamisawa, K. (2004) New method of denitrification analysis of *Bradyrhizobium* field isolates by gas chromatographic determination of ^15^N-labeled N_2_. Appl Environ Microbiol 70: 2886–2891.

Sánchez, C., Minamisawa, K. (2018) Redundant roles of *Bradyrhizobium oligotrophicum* Cu-type (NirK) and cd1-type (NirS) nitrite reductase genes under denitrifying conditions. FEMS Microbiol Lett 365: fny015.

Santos, M.S., Nogueira, M.A., Hungria, M. (2019) Microbial inoculants: reviewing the past, discussing the present and previewing an outstanding future for the use of beneficial bacteria in agriculture. AMB Expr 9: 205.

Schlesinger, W.H. (2009) On the fate of anthropogenic nitrogen. Proc Natl Acad Sci 106: 203–208.

Schwartz, W. (1972) Vincent, J.M, a manual for the practical study of the root-nodule bacteria (IBP Handbuch no. 15 des international biology program, London). XI u. 164 S., 10 Abb., 17 tab., 7 Taf. Oxford-Edinburgh 1970: Blackwell scientific Publ.,45 s. Z Allg Mikrobiol 12: 1521–4028.

Shapleigh, J.P. (2013) The prokaryotes: Prokaryotic physiology and biochemistry. In Rosenberg E, DeLong EF, Lory S, Stackebrandt E, Thompson F (eds). The prokaryotes. Springer: Berlin Heidelberg, pp 405–425.

Sistrom, W.R. (1962) The kinetics of the synthesis of photopigments in Rhodopseudomonas spheroides. J Gen Microbiol 28: 607–616.

Thompson, R.L., Lassaletta, L., Patra, P.K., Wilson, C., Wells, K.C., Gressent, A., et al. (2019) Acceleration of global N_2_O emissions seen from two decades of atmospheric inversion. Nat Clim Chang 9: 993–998.

Thomson, A.J., Giannopoulos, G., Pretty, J., Baggs, E.M., Richardson, D.J. (2012) Biological sources and sinks of nitrous oxide and strategies to mitigate emissions. Philos Trans R Soc B. 367: 1157–1168.

Thrall, P.H., Laine, A-L., Broadhurst, L.M., Bagnall, D.J., Brockwell, J. (2011) Symbiotic effectiveness of rhizobial mutualists varies in interactions with native Australian legume genera. PLoS ONE 6: e23545.

Tian, H., Xu, R., Canadell, J.G., Thompson, R.L., Winiwarter, W., Suntharalingam, P., et al. (2020) A comprehensive quantification of global nitrous oxide surces and sinks. Nature 586: 248–256.

Torres, M.J., Bueno, E., Jiménez-Leiva, A., Cabrera, J.J., Bedmar, E.J., Mesa, S., et al. (2017) FixK2 Is the Main Transcriptional Activator of *Bradyrhizobium diazoefficiens nosRZDYFLX* Genes in Response to Low Oxygen. Front Microbiol 8: 1621.

Torres, M.J., Simon, J., Rowley, G., Bedmar, E.J., Richardson, D.J., Gates, A.J., et al. (2016) Nitrous oxide metabolism in nitrate-reducing bacteria: physiology and regulatory mechanisms. In: Poole RK (eds). Advances in microbial physiology, first edition. Elsevier, pp 353–432.

Van Rossum, D., Muyotcha, A., Van Verseveld, H.W., Stouthamer, A.H., Boogerd, F.C. (1993) Effects of *Bradyrhizobium* strain and host genotype, nodule dry weight and leaf aera on groundnut *(Arachis hypogaea L. ssp. fastigiata)* yield. Plant and Soil 154: 279–288.

Van Rossum, D., Muyotcha, A., Van Verseveld, H.W., Stouthamer, A.H., Boogerd, F.C. (1994) Siderophore production by *Bradyrhizobium* spp. Strains nodulating groundnut. Plant and soil 163: 177–187.

Van Spanning, R., Richardson, D., Ferguson, S. (2007) Introduction to the biochemistry and molecular biology of denitrification. In: Bothe H,Ferguson SJ, Newton WE (eds). Biology of the nitrogen cylce. Elsevier: Amsterdam, The Netherlands, pp 3–21.

Velasco, L., Mesa, S., Delgado, M.J., Bedmar, E.J. (2001) Characterization of the *nirK* gene encoding the respiratory, Cu-containing nitrite reductase of *Bradyrhizobium japonicum*. Biochim Biophys Acta 1521: 130–134.

Velasco, L., Mesa, S., Xu, C., Delgado, M.J., Bedmar, E.J. (2004) Molecular characterization of *nosRZDFYLX* genes coding for denitrifying nitrous oxide reductase of *Bradyrhizobium japonicum*. Antonie Van Leeuwenhoek 85: 229–235.

Wang, R., Chang, Y.L., Zheng, W.T., Zhang, D., Zhang, X.X., Sui, X.H., et al. (2013) *Bradyrhizobium arachidis* sp. Nov., isolated from effective nodules of Arachis hypogaea grown in China. Syst Appl Microbiol 36: 101–105.

Winiwarter, W., Höglund-Isaksson, L., Klimont, Z., Schöpp, W., Amann, M. (2018) Technical opportunities to reduce global anthropogenic emissions of nitrous oxide. Environ Res Lett 13: 014011.

Woliy, K., Degefu, T., Frostegård, Å. (2019) Host Range and Symbiotic Effectiveness of N_2_O Reducing *Bradyrhizobium* Strains. Front Microbiol 10: 2746.

Xu, Z.F., Hansen, M.A., Hansen, L.H., Jacquiod, S., Sørensen, S.J. (2014) Bioinformatic Approaches Reveal Metagenomic Characterization of Soil Microbial Community. PLoS ONE 9: e93445.

Zhang, L., Wüst, A., Prasser, B., Müller, C., Einsle, O. (2019) Functional assembly of nitrous oxide reductase provides insights into copper site maturation. Proc Natl Acad Sci 116: 12822–12827.

Zhang, X.P., Nick, G., Kaijalainen, S., Terefework, Z., Paulin, L., Tighe, S.W., et al. (1999) Phylogeny and Diversity of *Bradyrhizobium Strains* isoalted from the Root Nodules of Peanut *(Arachis hypogaea)* in Sichuan, China. Syst Appl Microbiol 22: 378–386.

Zilli, .JÉ., Alves, B.J.R., Rouws, L.F.M., Simões-Araujo, J.L., de Barros Soares, L.H., Cassán, F., et al. (2019) The importance of denitrification performed by nitrogen-fixing bacteria used as inoculants in South America. Plant Soil 451: 5–24.

Zumft, W.G. (1997) Cell biology and molecular basis of denitrification. Microbiol Mol Biol Rev 61: 533–616.

