## Supplementary Figures and Tables_Gao et al for "Competition for electrons favors N_2_O reduction in denitrifying *Bradyrhizobium isolates*"

**Table S1.** Pairwise comparison of whole genome sequence similarities based on Average Nucleotide Identity (ANI). Strains that were highly similar according to the MLSA analysis of 4 housekeeping gene sequences are listed above the shaded area. For such strains, only one representative (marked in bold in this table) was included in the phylogenetic tree (Fig. 1). Some examples of comparisons of strains belonging to different clades, as determined by the MLSA analysis, are listed below the shaded area.

| Strain included in phylogenetic tree (Fig. 1) | Pairwise compared strain* | ANI estimate | Matches | Total |
|---|---|---|---|---|
| **HAMBI 2125** | HAMBI 2126 | **99.9830** | 2481 | 2584 |
| **HAMBI 2128** | HAMBI 2129 | **99.9919** | 3063 | 3106 |
| | HAMBI 2150 | **99.3635** | 2540 | 3106 |
| **HAMBI 2134** | HAMBI 2135 | **99.2192** | 2461 | 2803 |
| **HAMBI 2149** | HAMBI 2133 | **99.9839** | 2753 | 2820 |
| | HAMBI 2136 | **99.9507** | 2652 | 2820 |
| | HAMBI 2137 | **99.8860** | 2517 | 2820 |
| | HAMBI 2151 | **99.8258** | 2420 | 2820 |
| | HAMBI 2152 | **99.9122** | 2543 | 2820 |
| **HAMBI 2116** | HAMBI 2153 | 92.5675 | 1755 | 2508 |
| **HAMBI 2125** | HAMBI 2130 | 94.7212 | 2119 | 2584 |
| **HAMBI 2127** | HAMBI 2130 | 95.0357 | 2106 | 2551 |
| **HAMBI 2134** | HAMBI 2142 | 98.0163 | 2469 | 2803 |
| **HAMBI 2135** | HAMBI 2142 | 97.2603 | 2316 | 2695 |
| **HAMBI 2299** | HAMBI 2153 | 97.4109 | 2033 | 2489 |
| *B. diazoefficiens* USD110 | *B. diazoefficiens* SEMIA5080 | 98.7149 | 2802 | 3035 |

*Comparison with strain marked in bold to the left

**Table S2.** Denitrification gene operons of *Bradyrhizobium* strains, analyzed based on whole genome sequencing. Strains placed within the same section of the table were highly similar as seen from pairwise genome comparison, with ANI values >99.2, and in most cases >99.9 % (Table S1). They also had the same denitrification phenotype. Strains representing each of these clusters are marked in bold and were included in the phylogenetic tree (Fig. 1). Presence (+) or absence (-) of functional and regulatory/accessory genes are indicated. Colors represent a function detected after analysis of denitrification end products (blue= nitrate reduction; yellow = nitrite reduction; red = nitric oxide reduction; and green = nitrous oxide reduction). Blank = absence of function. The amino acid sequences of the *nor* operons of strains HAMBI 2299 and HAMBI 2153 were compared to those of the strains listed here (in bold) that produced functional reductases (except strain CCBAU 53363$^T$).

| *Bradyrhizobium* strains | Nitrate reduction | | | | | Nitrite reduction | | Nitric oxide reduction | | | | | Nitrous oxide reduction | | | | | | |
|---|---|---|---|---|---|---|---|---|---|---|---|---|---|---|---|---|---|---|---|
| | ***nap*** | | | | | ***nir*** | | ***nor*** | | | | | ***nos*** | | | | | | |
| | *E* | *D* | *A* | *B* | *C* | *K* | *V* | *C* | *B* | *Q* | *D* | *E* | *R* | *Z* | *D* | *Y* | *F* | *L* | *X* |
| **HAMBI 2125**<br>HAMBI 2126 | + | + | + | + | + | + | + | + | + | + | + | + | + | + | + | + | + | + | + |
| **HAMBI 2127** | + | + | + | + | + | + | + | + | + | + | + | + | + | + | + | + | + | + | + |
| **HAMBI 2130** | + | + | + | + | + | + | + | + | + | + | + | + | + | + | + | + | + | + | + |
| **HAMBI 3052$^T$** | + | + | + | + | + | + | + | + | + | + | + | + | + | + | + | + | + | + | + |
| ***CCBAU 53363$^T$** | + | + | + | + | + | + | + | + | + | + | + | + | + | + | + | + | + | + | + |
| **HAMBI 2115** | + | + | + | + | + | + | + | + | + | + | + | + | - | - | - | - | - | - | - |
| **HAMBI 2116** | + | + | + | + | + | + | + | + | + | + | + | + | - | - | - | - | - | - | - |
| **HAMBI 2128**<br>HAMBI 2129<br>HAMBI 2150 | + | + | + | + | + | + | + | + | + | + | + | + | - | - | - | - | - | - | - |
| **HAMBI 2134**<br>HAMBI 2135 | + | + | + | + | + | + | + | + | + | + | + | + | - | - | - | - | - | - | - |
| **HAMBI 2142** | + | + | + | + | + | + | + | + | + | + | + | + | - | - | - | - | - | - | - |
| ****HAMBI 2153** | + | + | + | + | + | + | + | + | - | - | + | + | - | - | - | - | - | - | - |
| **HAMBI 2299** | + | + | + | + | + | + | + | + | + | + | + | + | - | - | - | - | - | - | - |
| **HAMBI 2149**<br>HAMBI 2133<br>HAMBI 2136<br>HAMBI 2137<br>HAMBI 2151<br>HAMBI 2152 | + | + | + | + | + | + | + | - | - | - | - | - | - | - | - | - | - | - | - |

*The accession number for the complete genome sequence of CCBAU 53363 in NCBI is CP022219.

** A frame shift in *norB*; *norQ* is not in the correct reading frame.

**Table S3.** Genome accession numbers (JGI Gold study ID: Gs0134353).

| Genome Name / Sample Name | IMG Genome ID |
|---|---|
| *Bradyrhizobium sp.* HAMBI 2115 | 2824661429 |
| *Bradyrhizobium sp.* HAMBI 2116 | 2824679649 |
| *Bradyrhizobium sp.* HAMBI 2125 | 2824671348 |
| *Bradyrhizobium sp.* HAMBI 2126 | 2824687955 |
| *Bradyrhizobium sp.* HAMBI 2127 | 2824696289 |
| *Bradyrhizobium sp*. HAMBI 2128 | 2824753945 |
| *Bradyrhizobium sp*. HAMBI 2129 | 2824763712 |
| *Bradyrhizobium sp*. HAMBI 2130 | 2824773399 |
| *Bradyrhizobium sp*. HAMBI 2133 | 2824617872 |
| *Bradyrhizobium sp*. HAMBI 2134 | 2824609381 |
| *Bradyrhizobium sp.* HAMBI 2135 | 2824600985 |
| *Bradyrhizobium sp.* HAMBI 2136 | 2824635225 |
| *Bradyrhizobium sp*. HAMBI 2137 | 2824644064 |
| *Bradyrhizobium sp*. HAMBI 2142 | 2824653114 |
| *Bradyrhizobium sp*. HAMBI 2149 | 2824626560 |
| *Bradyrhizobium sp*. HAMBI 2150 | 2824704595 |
| *Bradyrhizobium sp*. HAMBI 2151 | 2824714736 |
| *Bradyrhizobium sp*. HAMBI 2152 | 2824723954 |
| *Bradyrhizobium sp*. HAMBI 2153 | 2824732956 |
| *Bradyrhizobium sp*. HAMBI 2299 | 2824746037 |
| *Bradyrhizobium sp*. HAMBI 3052 | 2731957961 |

**Table S4.** Primers and PCR conditions used in the current study.

| Nucleic acid | Primer (5'to 3') | Reference |
|---|---|---|
| 16S rRNA | 27F: AGA GTT TGA TCM TGG CTC AG<br>518R: ATT ACC GCG GCT GCT GG | (Weisburg et al., 1991; Muyzer *et al*., 1993) |
| *rpoB** | 454F: TCG CAG TTC ATG GAC CAG R<br>1364R: GTA GCC GTT CCA SGG CAT G | (Vinuesa *et al*., 2008; Aserse *et al*., 2012) |
| *atpD*** | 189-208: TCT GGT CCG YGG CCA GGA AG<br>804-784: CGA CAC TTC CGA RCC SGC CTG | (Stepkowski *et al*., 2005) |
| *recA**** | 41F: TTC GGC AAG GGM TCG RTS ATG<br>640R: ACA TSA CRC CGA TCT TCA TGC | (Vinuesa *et al*., 2005) |

* The annealing temperature for *rpoB* gene in the present study was 68 °C.

** The annealing temperature for *atpD* gene in the present study was 63 °C.

*** The annealing temperature for *recA* gene in the present study was 61 °C.

Weisburg, W.G., Barns, S.M., Pelletier, D.A., Lane, D.J. (1991) 16S Ribosomal DNA Amplification for Phylogenetic Study *J Bacteriol* **173**: 697–703.

Muyzer, G., De Waal, E.C., Uitterlinden, A.G. (1993) Profiling of Complex Microbial Populations by Denaturing Gradient Gel Electrophoresis Analysis of Polymerase Chain Reaction-Amplified Genes Coding for 16S rRNA. *Appl Environ Microbiol* **59**: 695–700.

Vinuesa, P., Rojas-Jiménez, K., Contreras-Moreira, B., Mahna, S.K., Prasad, B.N., Moe, H., *et al*. (2008) Multilocus Sequence Analysis for Assessment of the Biogeography and Evolutionary Genetics of Four *Bradyrhizobium* Species That Nodulate Soybeans on the Asiatic Continent. *Appl Environ Microbiol* **74**: 6987–6996.

Aserse, A.A., Räsänen, L.A., Aseffa, F., Hailemariam, A., Lindström, K. (2012) Phylogenetically diverse groups of *Bradyrhizobium* isolated from nodules of *Crotalaria* sp., *Indigofera* sp., *Erythrina brucei* and *Glycine max* growing in Ethiopia. *Mol Phylogenet Evol* **65**: 595–609.

Stepkowski, T., Moulin, L., Krzyzanska, A., McIlnnes, A., Law, I.J., John, H. (2005) European Origin of *Bradyrhizobium* Population Infecting Lupins and Serradella in Soil of Western Australia and South Africa. *Appl Environ Microbiol* **71**: 7041–7052.

Vinuesa, P., Silva, C., Werner, D., Martínez-Romero, E. (2005) Population genetics and phylogenetic inference in bacterial molecular systematics: The roles of migration and recombination in *Bradyrhizobium* species cohesion and delineation. *Mol Phylogenet Evol* **34**: 29–54.

**Table S5.** Accession numbers in NCBI for the genes used in Fig. 1.

| Species | *atpD* | *glnII* | *recA* | *rpoB* |
|---|---|---|---|---|
| *Bradyrhizobium* sp. AC70c | **MT319095** | MK631831 | **MT329654** | **MT329666** |
| *Bradyrhizobium* sp. AC86d2 | **MT319096** | MK631837 | MK631801 | **MT329667** |
| *Bradyrhizobium* sp. AC101b | **MT319097** | MK631850 | MK631815 | **MT329668** |
| *Bradyrhizobium* sp. AC87j1 | PTFE00000000 | PTFE00000000 | PTFE00000000 | PTFE00000000 |
| *B. algeriense* LMG 27618$^T$ | PYCM00000000 | PYCM00000000 | PYCM00000000 | PYCM00000000 |
| *B. americanum* CMVU44$^T$ | KC247125.1 | KX012942.1 | KC247141.1 | |
| *B. amphicarpaeae* HAMBI 3680$^T$ | CP029426 | CP029426 | CP029426 | CP029426 |
| *B. arachidis* CCBAU 051107$^T$ | HM107217.1 | HM107251.1 | HM107233.1 | JX437682.1 |
| *B. arachidis* CCBAU 23155 | GU433475.1 | GU433500.1 | GU433524 | JX437678.1 |
| *B. arachidis* CCBAU 33067 | GU433479.1 | GU433503.1 | GU433528.1 | JX437680.1 |
| *B. arachidis* CCBAU 45332 | JQ011347.1 | JQ011353.1 | JQ011350.1 | JX437681.1 |
| *B. betae* PL7HG1$^T$ | FM253129.1 | FJ970431.1 | FJ970378.1 | GU562860.1 |
| "*B. brasilense*" LMG 29353$^T$ | MPVQ00000000 | MPVQ00000000 | MPVQ00000000 | MPVQ00000000 |
| *B. cajani* AMBPC1010$^T$ | | KY349442.1 | KY349440.1 | WQNE01000000 |
| *B. canariense* BTA-1$^T$ | FM253135.1 | AY386765.1 | AY591553.1 | FM253263.1 |
| *B. centrosemae* A9$^T$ | KC247129.1 | KX012940.1 | KC247145.1 | |
| *B. cytisi* CTAW1$^T$ | LM994389.1 | GU001594.1 | GU001575.1 | LM994166.1 |
| *B. daqingense* CCBAU 15774$^T$ | HQ231289.1 | HQ231301.1 | KF962708.1 | JX437676.1 |
| *B. denitrificans* LMG 8443$^T$ | FM253153 | HM047121.1 | EU665419.1 | FM253282 |
| *B. diazoefficiens* USDA 110T | BA000040.2 | BA000040.2 | BA000040.2 | BA000040.2 |
| *B. elkanii* USDA 76T | KB900701 | KB900701 | AY591568.1 | KB900701 |
| *B. embrapense* SEMIA 6208T | HQ634875 | GQ160500.1 | HQ634899.1 | HQ634910 |
| *B. erythrophlei* CCBAU 53325T | | KF114693.1 | KF114669.1 | MG811654.1 |
| *B. ferriligni* CCBAU 51502T | | KJ818099.1 | KJ818112.1 | MG811655.1 |
| "*B. forestalis*" INPA54BT | PGVG00000000 | PGVG00000000 | PGVG00000000 | PGVG00000000 |
| *B. frederickii* USDA 10052T | SPQS00000000 | SPQS00000000 | SPQS00000000 | SPQS00000000 |
| *B. ganzhouense* RITF806T | JX277182.1 | JX277110.1 | JX277144.1 | |
| *B. guangdongense* CCBAU 51649T | KC508916.1 | KC509023.1 | KC509269.1 | KC509318.1 |
| *B. guangdongense* CCBAU 51658 | KC508918.1 | KC509025.1 | KC509271.1 | KC509320.1 |
| *B. guangdongense* CCBAU 51658 | KC508918.1 | KC509025.1 | KC509271.1 | KC509320.1 |
| *B. guangdongense* CCBAU 51670 | KC508902.1 | KC509008.1 | KC509254.1 | KC509303.1 |

Table S5. Cont.

| Species | *atpD* | *glnII* | *recA* | *rpoB* |
|---|---|---|---|---|
| *B. guangxiense* CCBAU 53363$^T$ | KC508926.1 | KC509033.1 | KC509279.1 | KC509328.1 |
| *B. guangxiense* CCBAU 53344 | KC508924.1 | KC509031.1 | KC509277.1 | KC509326.1 |
| *B. guangxiense* CCBAU 53429 | KC508931.1 | KC509038.1 | KC509284.1 | KC509333.1 |
| *B. huanghuaihaiense* CCBAU 23303$^T$ | LM994392.1 | HQ231639.1 | LM994321.1 | HQ428068.1 |
| *B. icense* HAMBI 3584$^T$ | KF896192.1 | KF896175.1 | JX943615.1 | CP016428.1 |
| *B. ingae* BR 10250$^T$ | KY753593.1 | KF927067.1 | KF927061.1 | KF927073.1 |
| *B. iriomotense*_NBRC_102520$^T$ | AB300994.1 | AB300995.1 | AB300996.1 | LM994170.1 |
| *B. japonicum* USDA 6$^T$ | AM168320.1 | AP012206.1 | AP012206.1 | AP012206.1 |
| *B. jicamae* PAC 68$^T$ | FJ428211.1 | FJ428204.1 | LM994324.1 | LM994173.1 |
| *B. kavangense* 14-3$^T$ | KY753592.1 | KM378446.1 | KM378399.1 | KM378311.1 |
| *B. liaoningense* LMG 18230$^T$ | FM253137.1 | AY386775.1 | FM253180.1 | FM253266.1 |
| *B. lupini* USDA 3051$^T$ | KU738808.1 | KM114862.1 | KM114866.1 | |
| "*B. macuxiense*"_HAMBI 3602$^T$ | LNCU00000000 | LNCU00000000 | LNCU00000000 | LNCU00000000 |
| *B. manausense* BR3351$^T$ | NZ_LJYG00000000 | KF785987.1 | KF785992.1 | KF785998.1 |
| *B. mercantei* LMG 30031$^T$ | NZ_MKFI01000006.1 | KX690621.1 | >KX690615.1 | NZ_MKFI01000001.1 |
| *B. namibiense* 5-10$^T$ | KX661387.1 | KM378440.1 | KM378377.1 | KM378306.1 |
| *B. neotropicale* BR10247$^T$ | LSEF01000046.1 | KJ661700.1 | KJ661714.1 | KF983829.2 |
| *B. niftali* CL 40$^T$ | SPQT00000000 | SPQT00000000 | SPQT00000000 | SPQT00000000 |
| "*B. nitroreducens*" KCTC 62391$^T$ | LFJC00000000 | LFJC00000000 | LFJC00000000 | LFJC00000000 |
| *B. oligotrophicum* LMG 10732$^T$ | JQ619232.1 | JQ619233.1 | JQ619231.1 | KF962713.1 |
| *B. ottawaense* OO99$^T$ | HQ455212.1 | HQ587750.1 | HQ587287.1 | HQ587518.1 |
| *B. pachyrhizi* PAC 48$^T$ | FJ428208.1 | FJ428201.1 | HM590777.1 | LM994172.1 |
| *B. paxllaeri* LMTR 21$^T$ | KF896186.1 | KF896169.1 | JX943617.1 | KP308154.1 |
| *B. retamae* Ro 19$^T$ | KC247101.1 | KC247108.1 | KC247094.1 | KF962714.1 |
| *B. rifense* CTAW71$^T$ | GU001617.1 | GU001604.1 | GU001585.1 | KC569468.1 |
| *B. ripae* LMG 30283$^T$ | | MF593086 | MF593090 | MF593098 |
| “*B. sacchari*"_HAMBI 3667$^T$ | KX065107 | KX065099 | KX065095 | NZ_LWIG01000014 |
| *B. shewense* ERR11T | NZ_FMAI01000019.1 | FMAI01000006.1 | NZ_FMAI01000022.1 | NZ_FMAI01000007.1 |
| *B. stylosanthis* BR 446T | LVEM01000002.1 | KU724148.1 | KU724163.1 | KU724166.1 |
| *B. subterraneum* 58 2-1T | KX661391.1 | KM378484.1 | KM378397.1 | KM378349.1 |

Table S5. Cont.

| Species | *atpD* | *glnII* | *recA* | *rpoB* |
|---|---|---|---|---|
| *B. symbiodeficiens* HAMBI 3684$^T$ | KP768551 | KP768609 | KF615036 | KP768667 |
| *B. tropiciagri* SEMIA 6148$^T$ | FJ390968.1 | FJ391048.1 | FJ391168.1 | HQ634909.1 |
| "*B. valentinum*" LMG 27619$^T$ | JX518561 | JX518575 | JX518589 | NZ_LLXX01000029 |
| *B. vignae* LMG 7-2$^T$ | | KM378443.1 | KM378374.1 | KM378308.1 |
| *B. viridifuturi* SEMIA 690$^T$ | NZ_LGTB01000039.1 | KR149131.1 | KR149140.1 | KU724169.1 |
| *B. yuanmingense* CCBAU 10071$^T$ | AY386760.1 | AY386780.1 | AY591566.1 | EF190174.1 |
| *Neorhizobium galegae* 540$^T$ | KF206641 | KF206809 | KF206896 | KF206983 |

The genus name *Bradyrhizobium* is abbreviated as *B.*, and the species names that have not been validate are shown in "". The accession numbers of the new data are shown in bold.

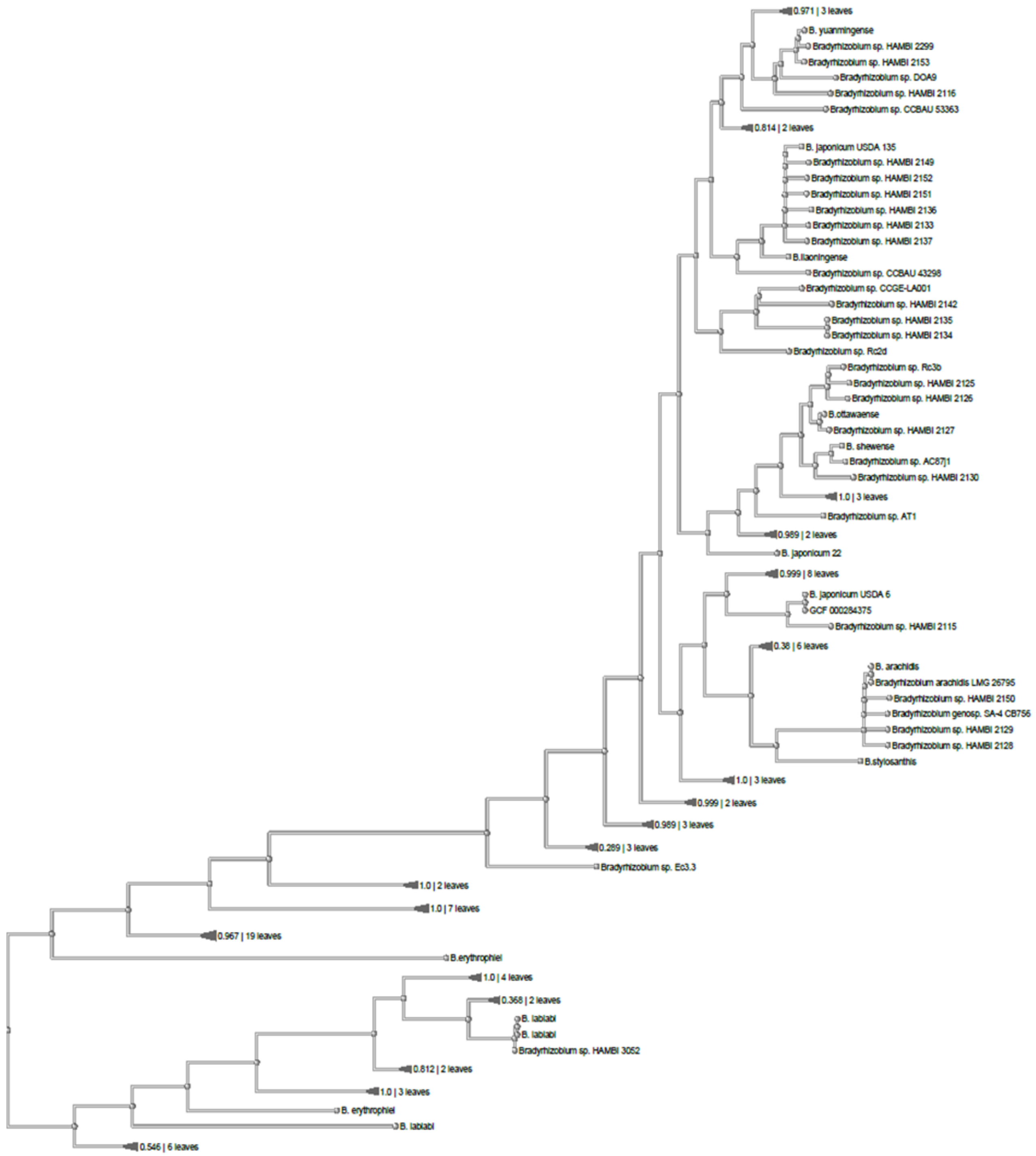

**Fig. S1.** Phylogenetic tree including all the HAMBI test strains listed in Table S2. The phylogenetic analysis was done by using the Genome Taxonomy Database Toolkit (GTDB-Tk) v0.3.2, which is based on a phylogeny inferred from the concatenation of 120 ubiquitous, single-copy proteins (Chaumeil *et al.*, 2020). The tree was drawn using the NCBI Genome Workbench (https://www.ncbi.nlm.nih.gov/tools/gbench).

Chaumeil, P.A., Mussig, A.J., Hugenholtz, P., Parks, D.H. (2020) GTDB-Tk: a toolkit to classify genomes with the Genome Taxonomy Database. *Bioinformatics* **36**: 1925-1927.

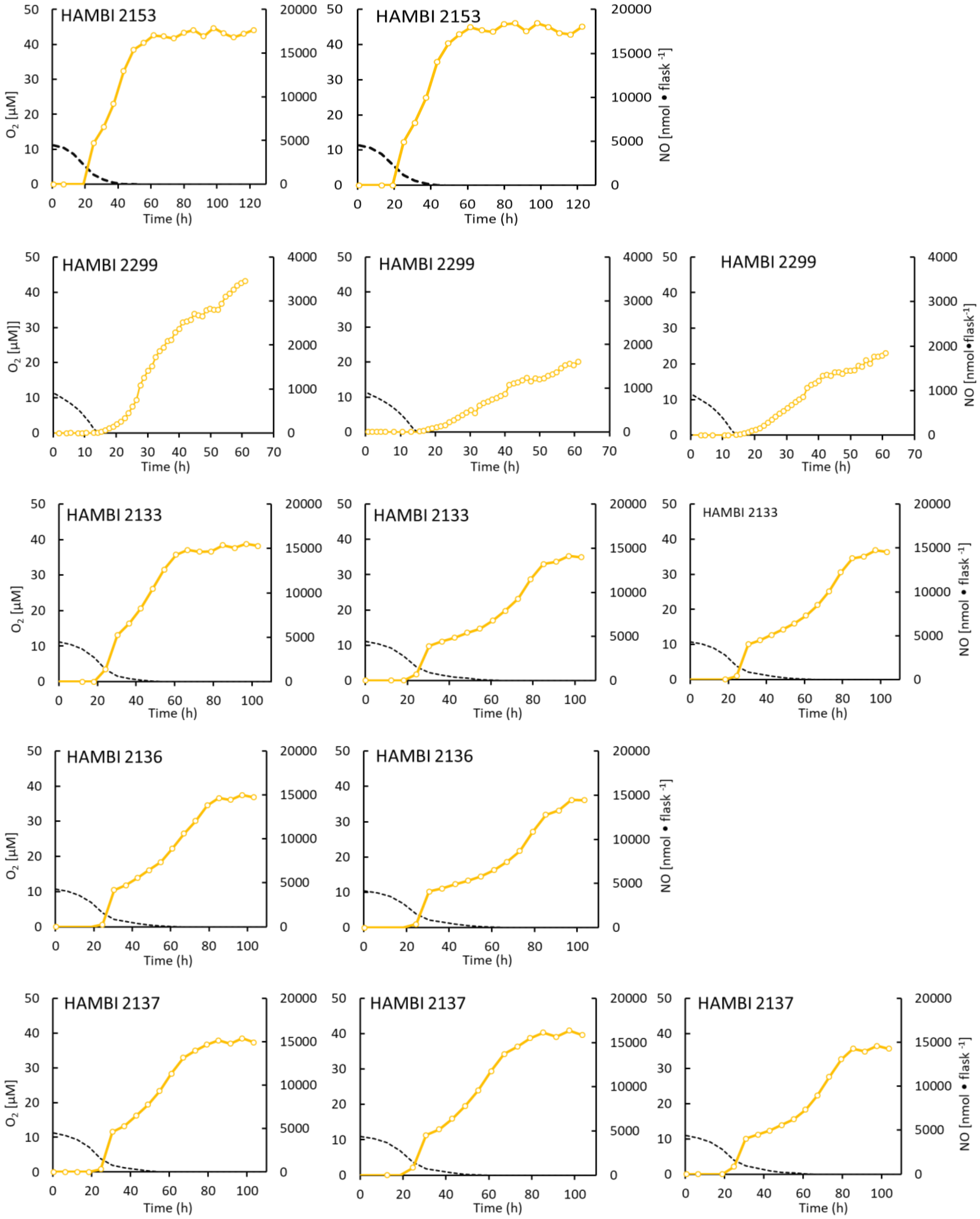
HAMBI 2153
HAMBI 2153
HAMBI 2299
HAMBI 2299
HAMBI 2299
HAMBI 2133
HAMBI 2133
HAMBI 2133
HAMBI 2136
HAMBI 2136
HAMBI 2137
HAMBI 2137
HAMBI 2137
$O_2$ [µM]
NO [nmol • flask$^{-1}$]
Time (h)

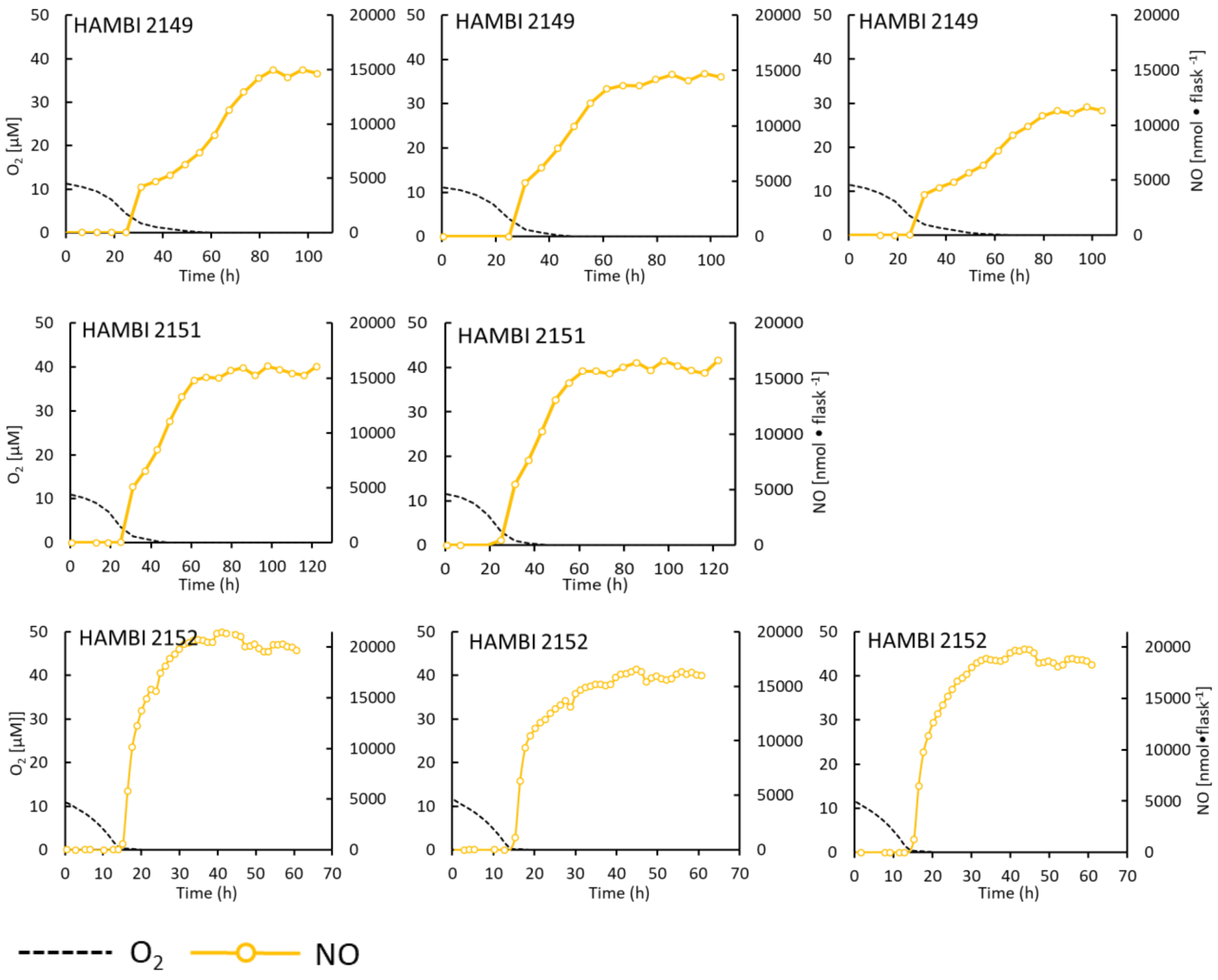

**Fig. S2.** Gas kinetics during and after transition from aerobic to anaerobic respiration for the strains apparently unable to reduce NO (results from 2 or three replicate flasks are shown). The cultures were incubated in YMB medium supplemented with 1 mM $KNO_3$ and with 0.7 ml $O_2$ (around 10 µM $O_2$ in liquid) and 1 ml $N_2O$ (8000 ppmv) in the headspace. The average rates of $N_2O$ production were as low as 6-12 nmol flask$^{-1}$ h$^{-1}$ (average of the replicates throughout the entire experiment), which is not significantly different from zero.

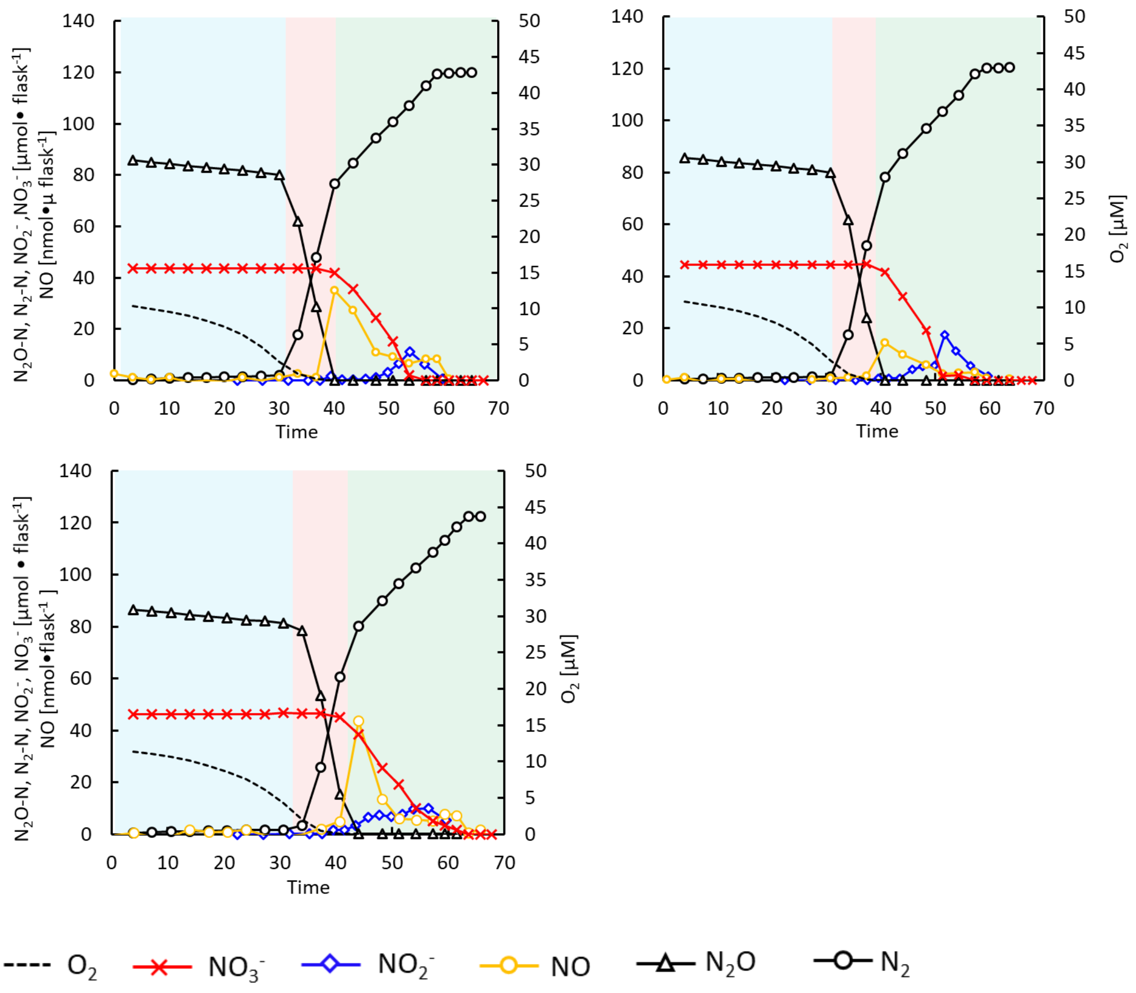

**Fig. S3.** Gas- and $NO_2^-$ kinetics for *Bradyrhizobium* strain HAMBI 2125 during and after transition from aerobic to anaerobic condition. The cultures (four replicate flasks , one shown in Fig. 2A, main paper) were grown in YMB supplemented with 1 mM $KNO_3$ and with 1 % $O_2$ (approximately 10 µM $O_2$ in liquid) and 1.2 ml $N_2O$ in the headspace (approximately 90 µmol $N_2O$-N flask$^{-1}$, corresponding to 0.26 mM $N_2O$ in the liquid). There was a marginal $NO_2^-$ accumulation before depletion of exogenous $N_2O$. During the subsequent reduction of $NO_3^-$, there was a transient accumulated 10 to 20 µmol $NO_2^-$ flask$^{-1}$, accounting for 20-40 % of the initial $NO_3^-$-N (50 µmol $NO_3^-$-N via$^{-1}$). Background colors indicate the dominating respiratory activity. Blue: $O_2$ respiration; Pink: $N_2O$ respiration; Green: Complete denitrification ($NO_3^-$ to $N_2$).

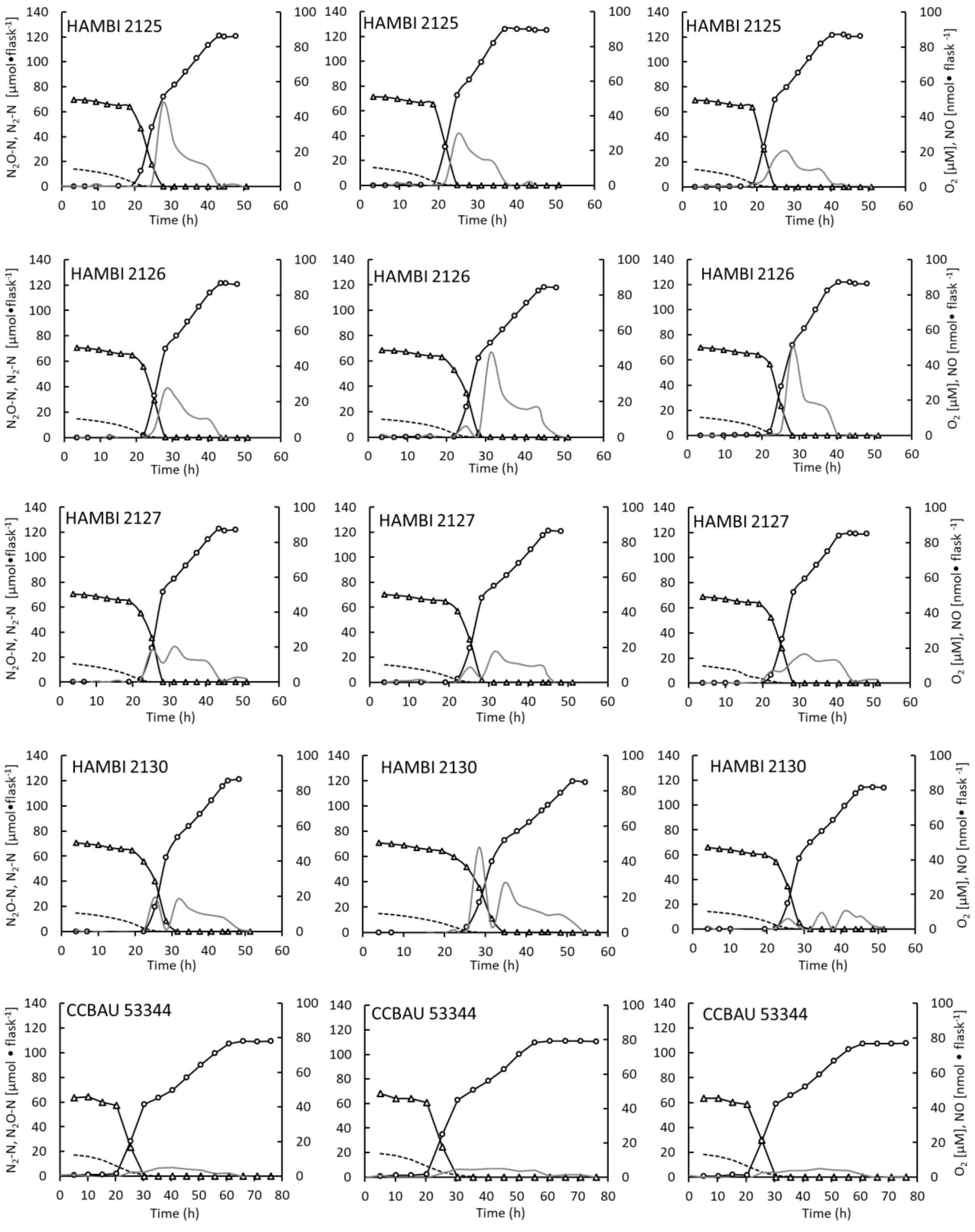
HAMBI 2125
HAMBI 2125
HAMBI 2125
HAMBI 2126
HAMBI 2126
HAMBI 2126
HAMBI 2127
HAMBI 2127
HAMBI 2127
HAMBI 2130
HAMBI 2130
HAMBI 2130
CCBAU 53344
CCBAU 53344
CCBAU 53344
N2O-N, N2-N [µmol•flask-1]
N2-N, N2O-N [µmol • flask-1]
O2 [µM], NO [nmol• flask -1]
Time (h)

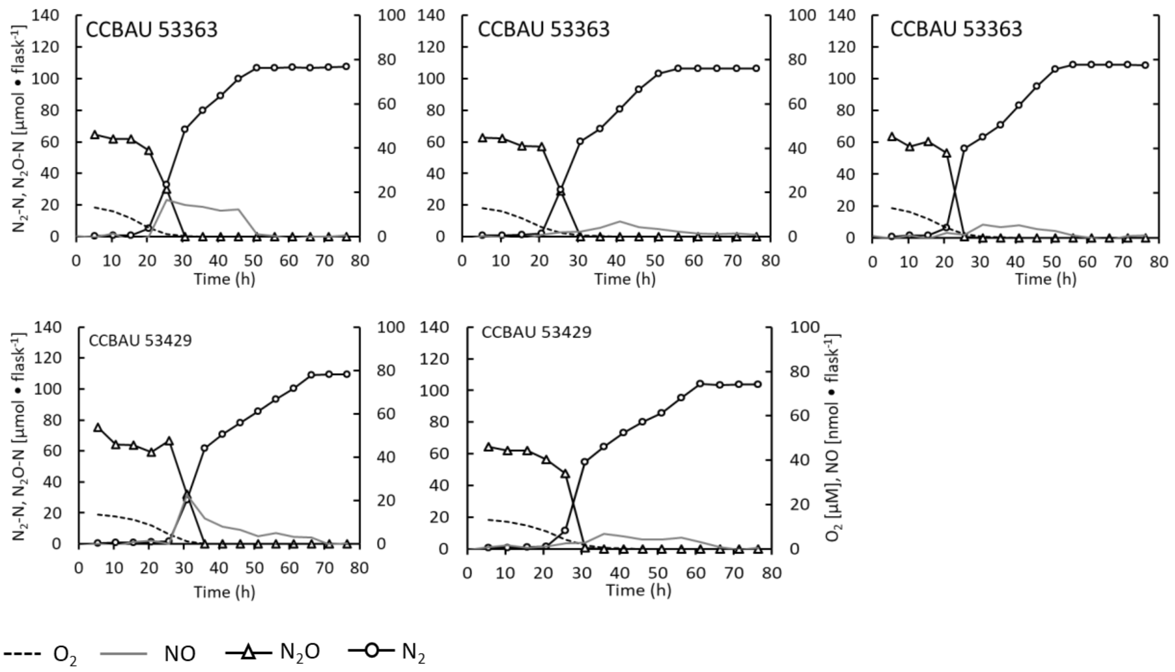

**Fig. S4.** Gas kinetics for the $N_2O$-reducing *Bradyrhizobium* strains (three replicate flasks except for CCBAU 53429) during and after transition from aerobic to anaerobic conditions. The cultures were incubated in YMB medium supplemented with 1 mM $KNO_3$ and with 0.7 ml $O_2$ (approximately 10 µM $O_2$ in liquid) and ~1 ml initial $N_2O$ in the headspace. All cultures showed the same strong preference for $N_2O$ over $NO_3^-$ in the presence of exogenous $N_2O$. This phase was followed by a phase where $N_2$ was produced from reduction of the reduction of $NO_3^-$. The $N_2$ plateaus were 110-140 µmol $N_2$-N flask$^{-1}$, depending on the exact amount of $N_2O$ that had been added, which corresponds to near 100 % recovery of $NO_3^-$ + $N_2O$-N as $N_2$ in all flasks.

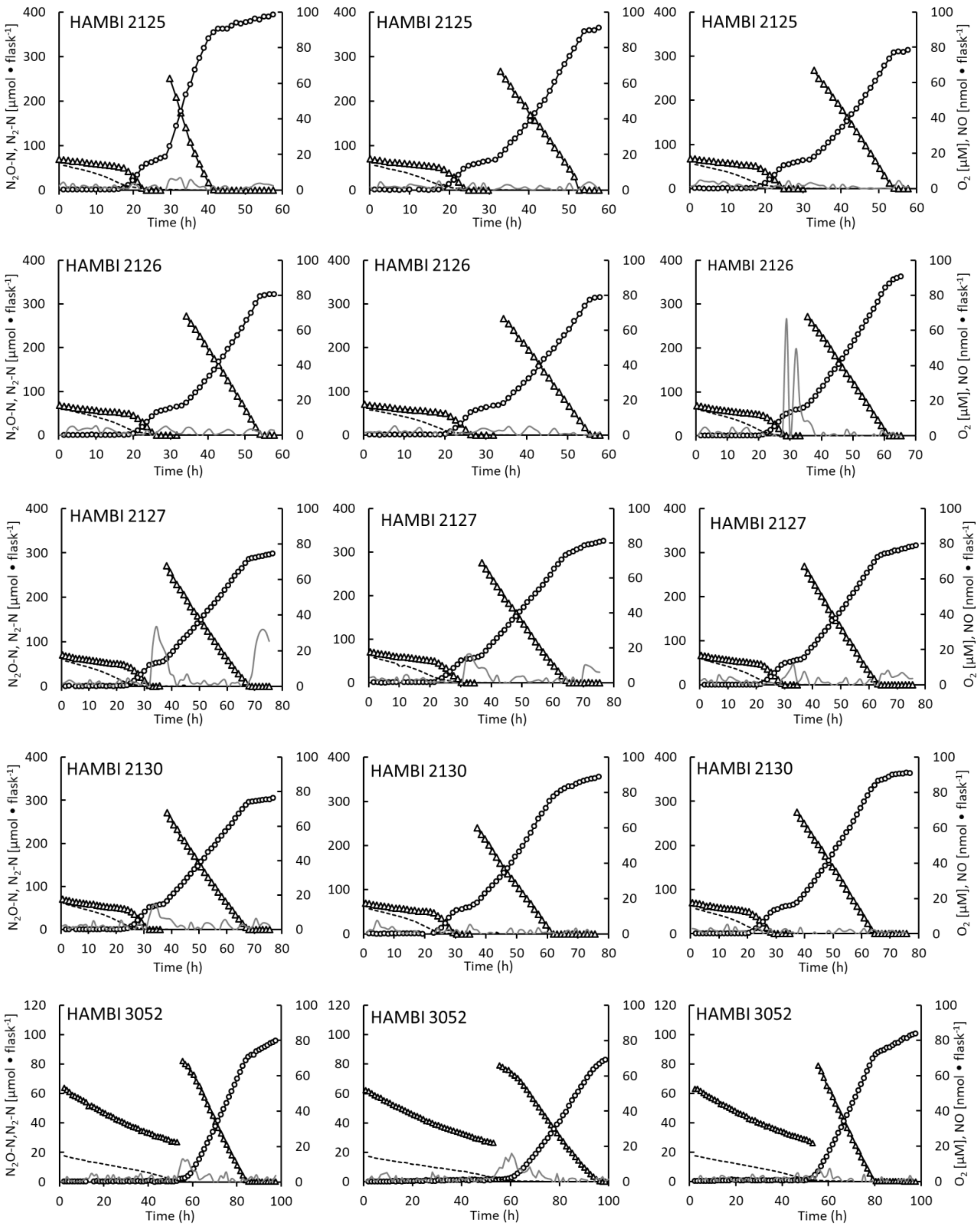
HAMBI 2125
HAMBI 2126
HAMBI 2127
HAMBI 2130
HAMBI 3052
N$_2$O-N, N$_2$-N [µmol • flask$^{-1}$]
O$_2$ [µM], NO [nmol • flask$^{-1}$]
Time (h)

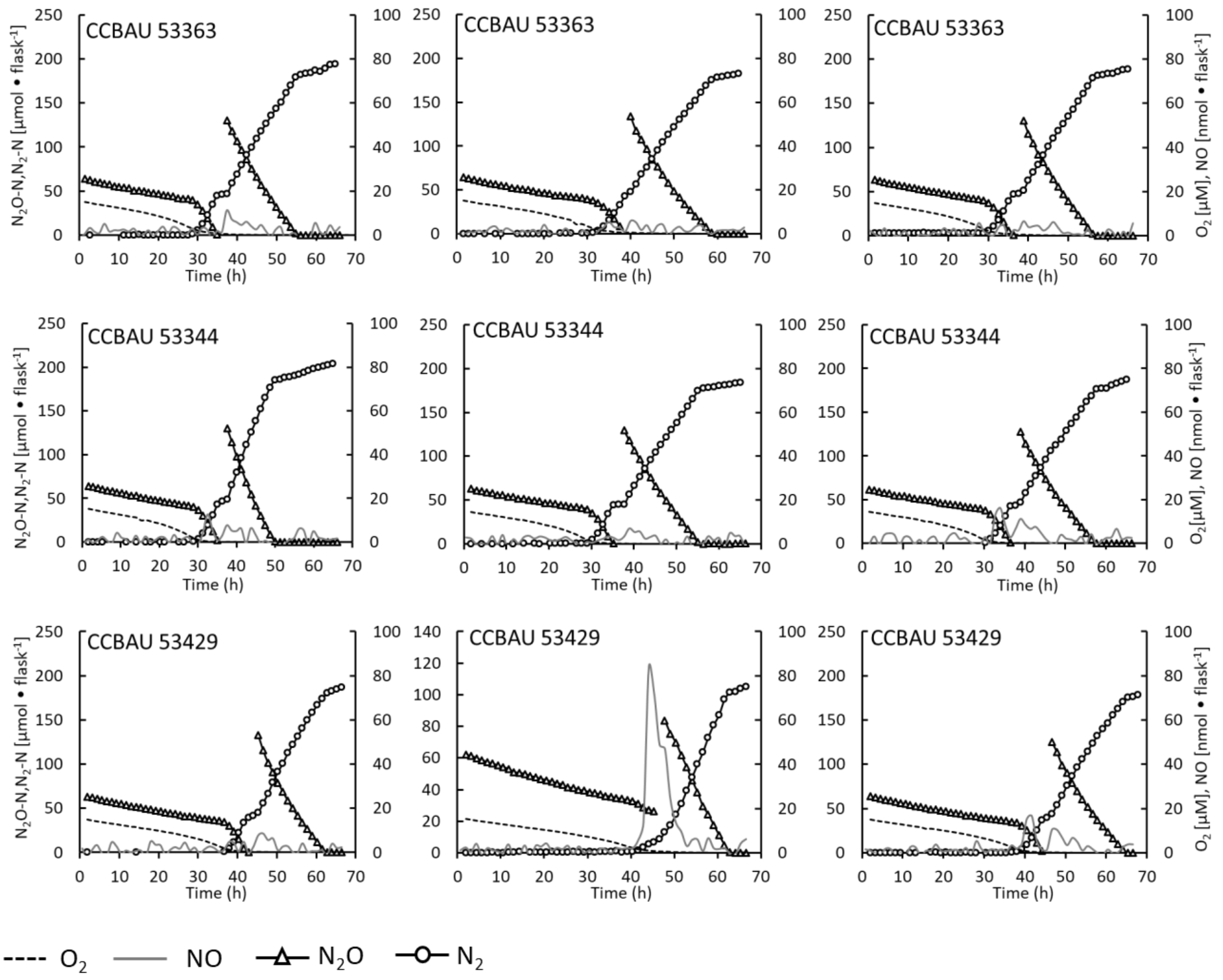

**Fig. S5.** Gas kinetics for the $N_2O$-reducing *Bradyrhizobium* strains during and after transition from aerobic to anaerobic conditions (three replicate flasks). The cultures were incubated in YMB medium supplemented with 2 or 3 mM $KNO_3$ in the medium, and 1 ml $O_2$ (around 15 µM $O_2$ in liquid) and 1 ml $N_2O$ (around 70-80 µmol $N_2O$-N) in the headspace. A second injection of $N_2O$ (around 80 or 150 or 260 µmol $N_2O$-N) was done when the cultures were actively denitrifying $NO_3^-$ to $N_2$. The cultures preferentially reduced the exogenously provided $N_2O$ over reducing $NO_3^-$, seen from the immediate hampering of $NO_3^-$ reduction by $N_2O$ and the lower rate of $N_2$-production from $NO_3^-$ reduction compared to that from reduction of exogenous $N_2O$.

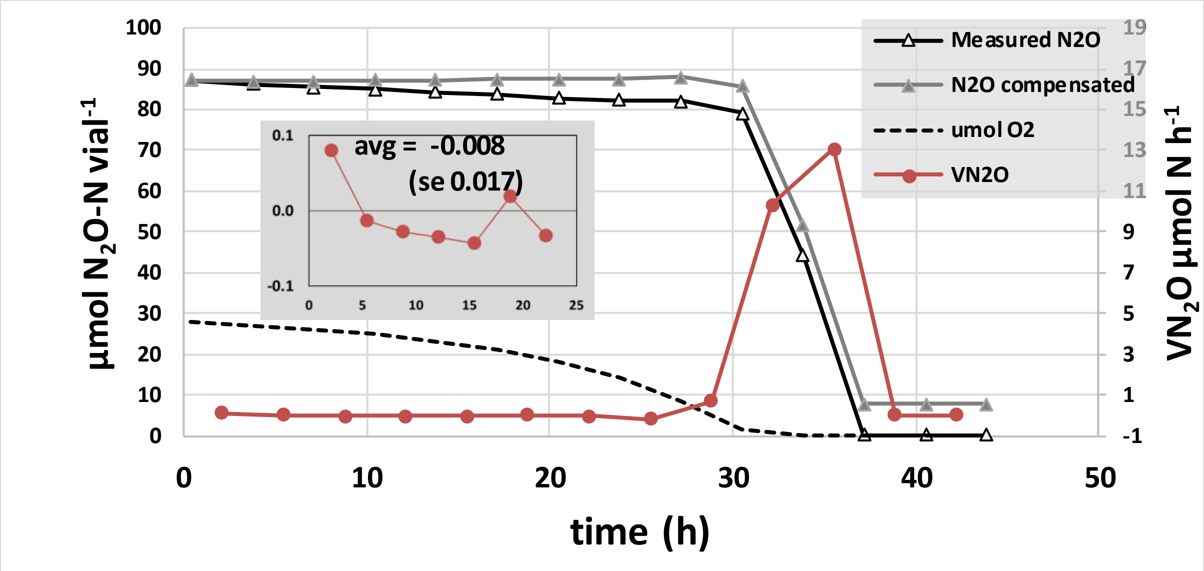

**Fig. S6.** Correction for sampling loss. Gradual decline in measured $N_2O$ is invariably observed during the oxic phase of the experiments (Fig S3-S5, Fig 2 and 4 in the main paper). This is not due to $N_2O$-reduction, but to sampling loss. Here we demonstrate the phenomenon for a single flask from the experiment shown in full in Fig 2. The panel shows measured $N_2O$ and $O_2$, the estimated rate of $N_2O$ reduction (which takes sampling loss into account), and an additional $N_2O$-curve which is the measured $N_2O$ + cumulated sampling loss ("$N_2O$ compensated"). The curve for "$N_2O$ compensated" does not decline during the oxic phase and does not reach zero for an obvious reason: the bacteria are unable to reduce the $N_2O$ that has been removed by sampling. This is why we present measured $N_2O$, rather than $N_2O$ compensated for sampling loss, in the figures in the paper. The inserted panel (grey) shows the estimated rate of $N_2O$ reduction during the oxic phase at higher resolution, seen to fluctuate around zero (average and standard error is shown in the panel). The sampling loss depends on the robot used: our new robots (Molstad *et al.*, 2007) take smaller samples than the original (Molstad *et al.*, 2016) but may also vary from one experiment to the other with the same robot due to wear of the peristaltic tube. Therefore, each experiment includes flasks with standard gases, which are used to estimate the actual sampling loss.

Molstad, L., Dörsch, P., Bakken, L.R. (2007) Robotized incubation system for monitoring gases ($O_2$, NO, $N_2O$ $N_2$) in denitrifying cultures. *J Microbiol Methods* **71**: 202–211.

Molstad, L., Dörsch, P., Bakken, L.R. (2016) Improved robotized incubation system for gas kinetics in batch cultures. *ResearchGate*. https://doi.org/10.13140/RG.2.2.30688.07680.

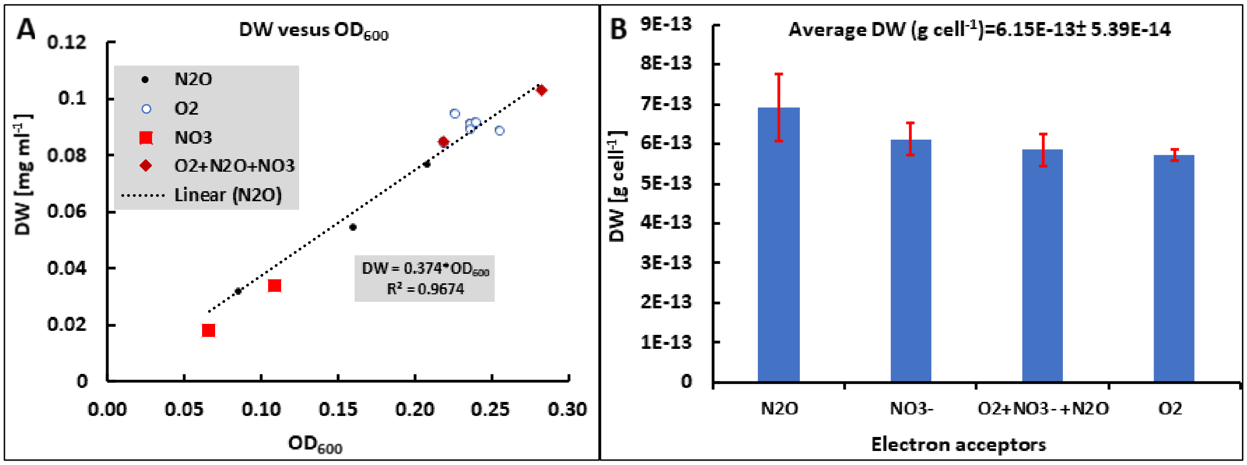

**Fig. S7.** Relationship between $OD_{600}$ and cell dry weight for *Bradyrhizobium* strain HAMBI 2125. Panel A. Measured cell dry weight (DW) plotted against measured $OD_{600}$. Each point is the endpoint value for single flasks with growth based on respiration of either $O_2$ (180 µmol flask$^{-1}$), $N_2O$ (800 µmol flask$^{-1}$), $NO_3^-$ (200 µmol flask$^{-1}$) or flasks with all three electron acceptors (marked $O_2$+$N_2O$+$NO_3$ in the legend). The regression is based on all data. Panel B. Calculated DW per cell based on microscopic counts and measured $OD_{600}$ of samples taken after 16 and 40 h of growth (n=3 for each treatment). The cell DW was calculated from the relationship between measured cell DW and $OD_{600}$ shown in panel A (mg DW ml$^{-1}$ = $OD_{600}$*0.374). The estimated cell DW was higher for cells grown with $N_2O$ as electron acceptor, but the difference was not statistically significant. Therefore, the average cell dry weight of 6.15E-13±5.39E-14 g cell$^{-1}$ (sd; n=12) was used for converting cell DW (based on OD) to cell density.

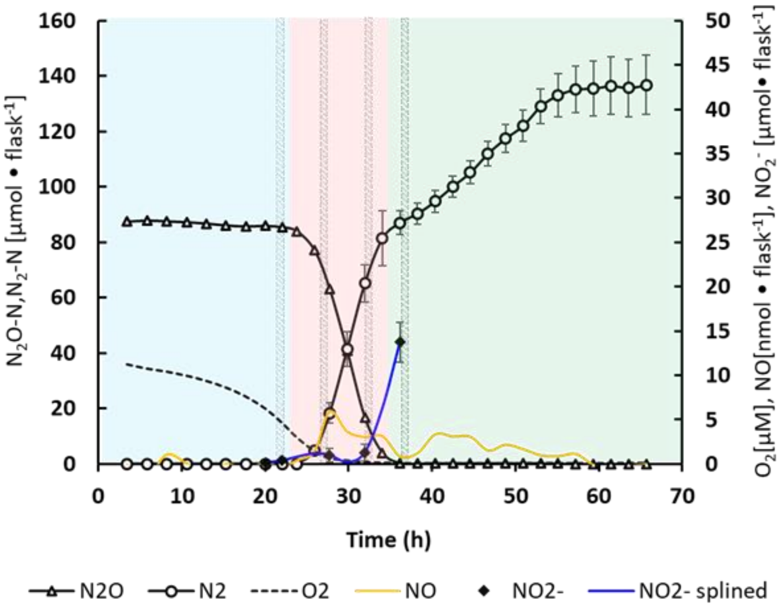

**Fig. S8.** Gas and $NO_2^-$ kinetics for *Bradyrhizobium* strain HAMBI 2125 in the incubation experiment used for proteomics. The cultures were incubated in 50 ml YMB medium supplied with 1 mM $KNO_3$ and 1% $O_2$ and 1 ml $N_2O$ (around 80 µmol N flask$^{-1}$) in the headspace. Cells were harvested for proteomics and $NO_2^-$ analyses at the time points indicated by gray bars (in 22, 27, 32 and 36 hpi). An additional $NO_2^-$ measurement was done at 20 hpi. Bars marked in the curves represent standard deviation (n=3 for each sampling).
